# The primate gut bacterial microbiome: a systematic review of research methodologies, taxonomic coverage, and conservation implications

**DOI:** 10.64898/2026.06.18.733207

**Authors:** Tommy Charles Burch, Paris Grace Badrock, Jean P Boubli, Naiara Guimarães Sales

## Abstract

Primates are central to both human evolutionary research and ecosystem functioning, serving as seed dispersers, predators, pollinators, and prey. Despite their value to human and ecosystem science, global primate populations continue to decline, with ∼65% of species currently threatened with extinction. Conservation biology increasingly recognises that survival depends not only on protecting habitats and populations, but also on safeguarding the microbial communities that underpin host health, nutrition, and resilience. The gut bacterial microbiome plays a critical role in digestion, immune function, and adaptation to environmental change, making it an important dimension of primate conservation. Here, we systematically and quantitatively assessed the taxonomic and geographic coverage of primate gut bacterial microbiome research to identify key knowledge gaps relevant to primate conservation. Between 2001 and 2025, 261 articles were published across 100 journals. While taxonomic coverage is high at the family level, it declines substantially at the finer taxonomic scales. Currently, ∼34.5% of species have been studied, leaving gut bacterial biodiversity undocumented for 344 species. Moreover, approximately one-third of studied species have exclusively been studied in captivity, limiting insights into natural microbiome variation and reducing the conservation relevance of these findings. Geographic biases further hinder conservation applications, with megadiverse countries such as Brazil, the Democratic Republic of Congo, and Indonesia underrepresented. In addition, study methodology and reporting standards remain inconsistent. To address these challenges, a framework for the standardised reporting of a minimum set of data for primate gut bacterial microbiome research is included in this review. Adoption of this framework will improve transparency, comparability, and data accessibility, thereby enhancing the utility of microbiome research for primate conservation. By integrating microbial ecology into conservation biology, we highlight the microbiome as a potential critical frontier for safeguarding primate health, evolutionary potential, and long-term survival.

## INTRODUCTION

Over the past century, non-human primates have been central to evolutionary and biomedical research due to their close similarity to humans in anatomy, physiology, cognition, reproduction, and their genetic relatedness (The Chimpanzee Sequencing and Analysis Consortium, 2005; Shively & Clarkson, 2009; Phillips et al., 2014; Picaud et al., 2019). Beyond their relevance for human evolution and biomedical research, primates are also the focus of extensive ecological research, as they play crucial roles–as seed dispersers (Koné et al., 2008; Sengupta et al., 2015; Razafindratsima et al., 2018), seed predators (Mourthé et al., 2008; Chen et al., 2023b), pollinators (Birkinshaw & Colquhoun, 1998; Heymann, 2011), and even prey (Zuberbühler & Jenny, 2002; Santos et al., 2014). These traits make primates essential for the maintenance of ecosystem health and underscores clear scientific and conservation-based rationale for studying primates.

However, ∼65% of primate species are currently threatened with extinction (Estrada & Garber, 2022; Fernández et al., 2022), and nearly one quarter of species whose conservation status has been re-assessed have declined (Wang et al., 2025). Although primates show great potential to adapt to meet ecological challenges over evolutionary timescales (Roos et al., 2025), many primates, particularly ecological specialists, are vulnerable to habitat loss, fragmentation, and human encroachment (Schwitzer et al., 2011). Human-driven ecosystem and resource alterations broaden the wildlife-human interface, increasing potential zoonotic and reverse-zoonotic spillover events (Patz et al., 2004; Al Norman et al., 2024; Boubli et al., 2026). The global decline in primate populations, together with the expansion of the wildlife-human interface, provides a strong rationale for studying primate gut microbiomes to better understand their natural state and how they are shaped by environmental *stimuli*, including those increasingly introduced through anthropogenic activities.

A microbiome can be described as a community of microorganisms inhabiting a distinct environment, such as the skin, genital, oral cavity, or gut. The gut microbiome refers to microbial communities inhabiting the gastrointestinal tract and are typically composed of bacteria, viruses, fungi, archaea, and protists, with bacteria usually the most abundant in humans and other animals (Qin et al., 2010; Henderson et al., 2015; Li et al., 2021). Study into primate gut microbiomes has grown in interest due to the insights that can be obtained relating to host health and disease status, diet and nutrition, behaviour, and life histories (Stumpf et al., 2016). Moreover, these samples can often be collected non-invasively, minimising the disturbance inadvertently caused by observational wildlife monitoring.

Several reviews have synthesised primate microbiome research from conservation, ecological, evolutionary, and comparative perspectives (Amato, 2016; Stumpf et al., 2016; Davenport et al., 2017; Clayton et al., 2018; Amato & Stumpf, 2019; Björk et al., 2019; Bambi et al., 2025). Some reviews have been thematic, for example emphasising comparative insights between humans, primates and other mammals (Davenport et al., 2017), temporal dynamics and the value of longitudinal sampling (Björk et al., 2019), and summarising multi-taxon studies to understand the dynamics and interactions within the gut microbiome of primates (Bambi et al., 2025). Additionally, some studies have provided an overview of methodologies used, benefits and challenges of available methods, and promoted the importance of the standardisation of protocols moving forward (Clayton et al., 2018; Amato & Stumpf, 2019).

Whilst previous reviews have provided valuable synthesis, especially by summarising key findings and delving into many specific host-microbe associations, none have quantified the taxonomic or geographic research effort. This work offers a distinct, quantitatively focused synthesis compared to earlier foundational or thematic reviews, highlighting biases and gaps in the current literature to guide future research using these methodological approaches to support primate conservation efforts. Additionally, this review adds to the discussion of methodologies used, by providing a timeline of the temporal trends related to publication frequency, methodologies used, and research scopes covered across the literature.

In this systematic review, all eligible scientific articles investigating the gut bacterial communities of wild and captive primates were compiled. From each article, data were extracted on primate taxonomy, geographic distribution, lifestyle type (wild or captive), and other key methodological variables. The dataset was used to evaluate taxonomic coverage and distribution of primate gut bacterial microbiome studies and to examine temporal trends in publication frequency, study scope, and methodologies used. Research biases and gaps were highlighted to support the applicability of these research approaches for primate conservation moving forward. Finally, this review proposes standardised reporting of a minimum set of data in primate microbiome studies to improve accessibility and transparency of data within published studies. The implementation of this framework can ultimately lead to greater comparability between studies and enhance the applicability of microbiome studies for primate conservation efforts.

## METHODS

### Library curation

Systematic searches were conducted on Web of Science (WoS) and Scopus to identify studies investigating the primate gut bacterial microbiome up to December 2025, with no restriction on the earliest publication date. A combination of keywords related to primates, microbiomes, and conservation was applied using Boolean operators (OR, AND) (Table SI). We followed the PRISMA 2020 guidance (Page et al., 2021) and adapted the flow diagram for this review (Figure 1). Identification, screening, eligibility, and inclusion numbers (and justification for exclusion) are summarised in Figure 1, with detailed information included in the Supplementary Material (Table SII). Search results retrieved 2,766 records (including books, chapters, notes, etc.). An initial filtering step was conducted, removing sources other than research articles (e.g., books and notes) with a total of 364 records excluded. The remaining 2,402 scientific articles were exported to EndNote, and duplicates were removed (184 articles). To ensure comprehensive coverage, we also conducted targeted non-systematic searches on Google Scholar and Google, which identified 52 additional publications.

**Figure 1:**
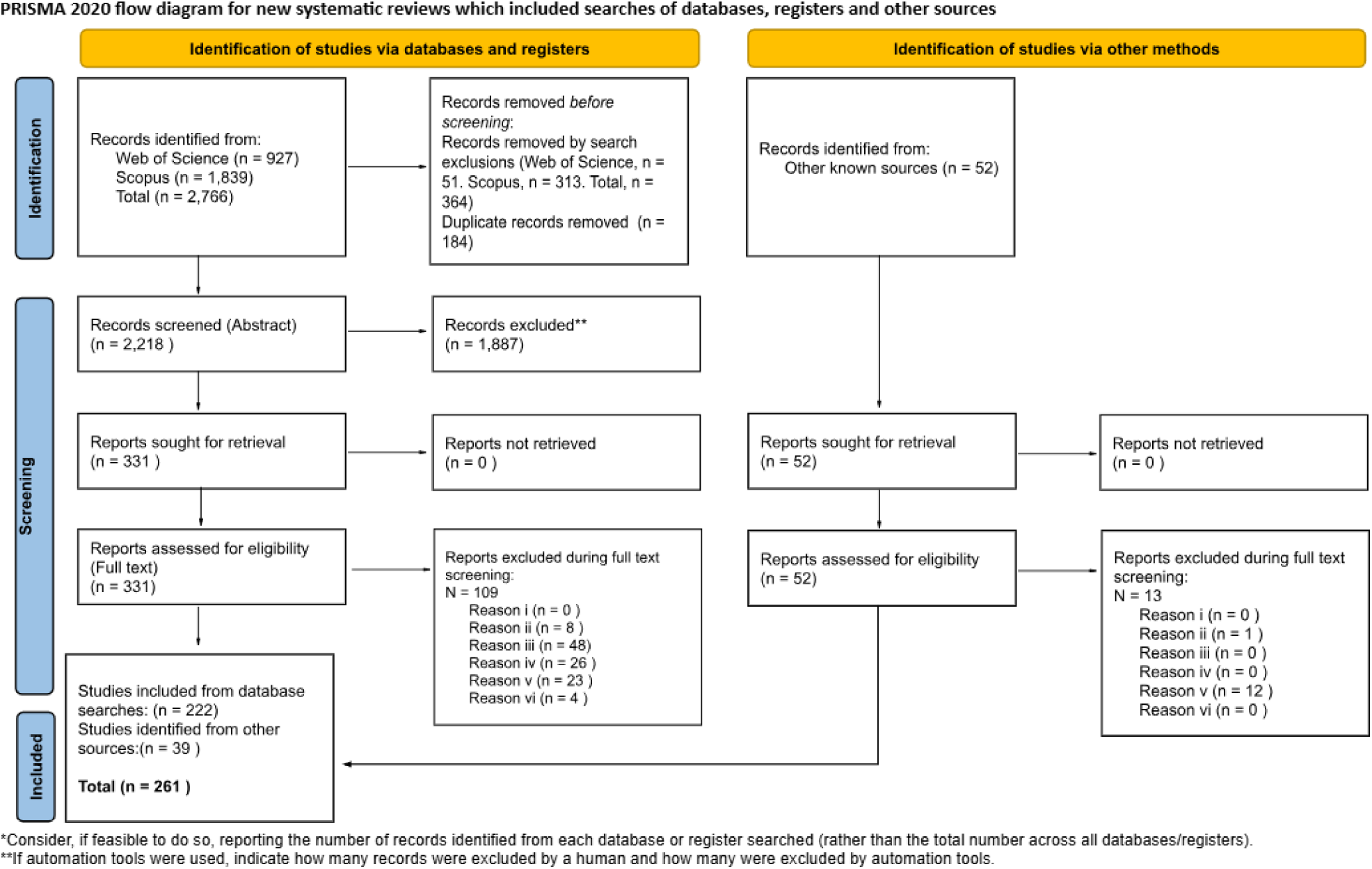
The number of studies included and excluded for this review. PRISMA flow diagram (Page et al. 2021), modified for this review. Licence: <u>Deed - Attribution 4.0 International - Creative Commons</u>

Abstracts were screened using eligibility criteria including six main components, considering (i) publication type, (ii) data type, (iii) microbial community focus, (iv) anatomical focus, (v) study setting and population, and (vi) language of publication (Table SII). After filtering, 331 of the 2,218 articles retrieved from WoS and Scopus were retained. The remaining articles were fully screened (i.e., considering all sections) for eligibility. Of the remaining 331 articles, 109 were excluded based on data type (8), microbial community focus (48), anatomical focus (26), study setting and population (23), and language of publication (4) criteria. The 52 articles that were identified via non-systematic search methods were then assessed via their main text, of which 39 met the eligibility criteria; exclusions were made based on data type (1) and study setting and population (12). In total, 261 scientific articles were included in this review for quantitative and narrative syntheses. The publications included in this review are available in Supplementary File 1 and those excluded are available in Supplementary File 4.

### Data analysis

From each article, 26 data points were extracted (Table SIII). Data extraction was conducted in Excel through a manual curation of research articles. All subsequent analyses were conducted in R (version 4.4.3; RStudio version 2026.04.0+526). For full data analysis, see Supplementary Material.

## RESULTS

### Publication frequency, distribution, and study scope

Between 2001 and 2025, 261 eligible articles spanning 100 journals were identified (Figure 2) (Table SV). Publications were concentrated in the American Journal of Primatology (n=31), Scientific Reports (n=20), and Frontiers in Microbiology (n=17). Since 2020, this distribution has shifted toward multidisciplinary outlets, led by Frontiers in Microbiology (n=16), followed by Scientific Reports (n=11), reflecting a broader disciplinary uptake of primate microbiome research.

**Figure 2:**
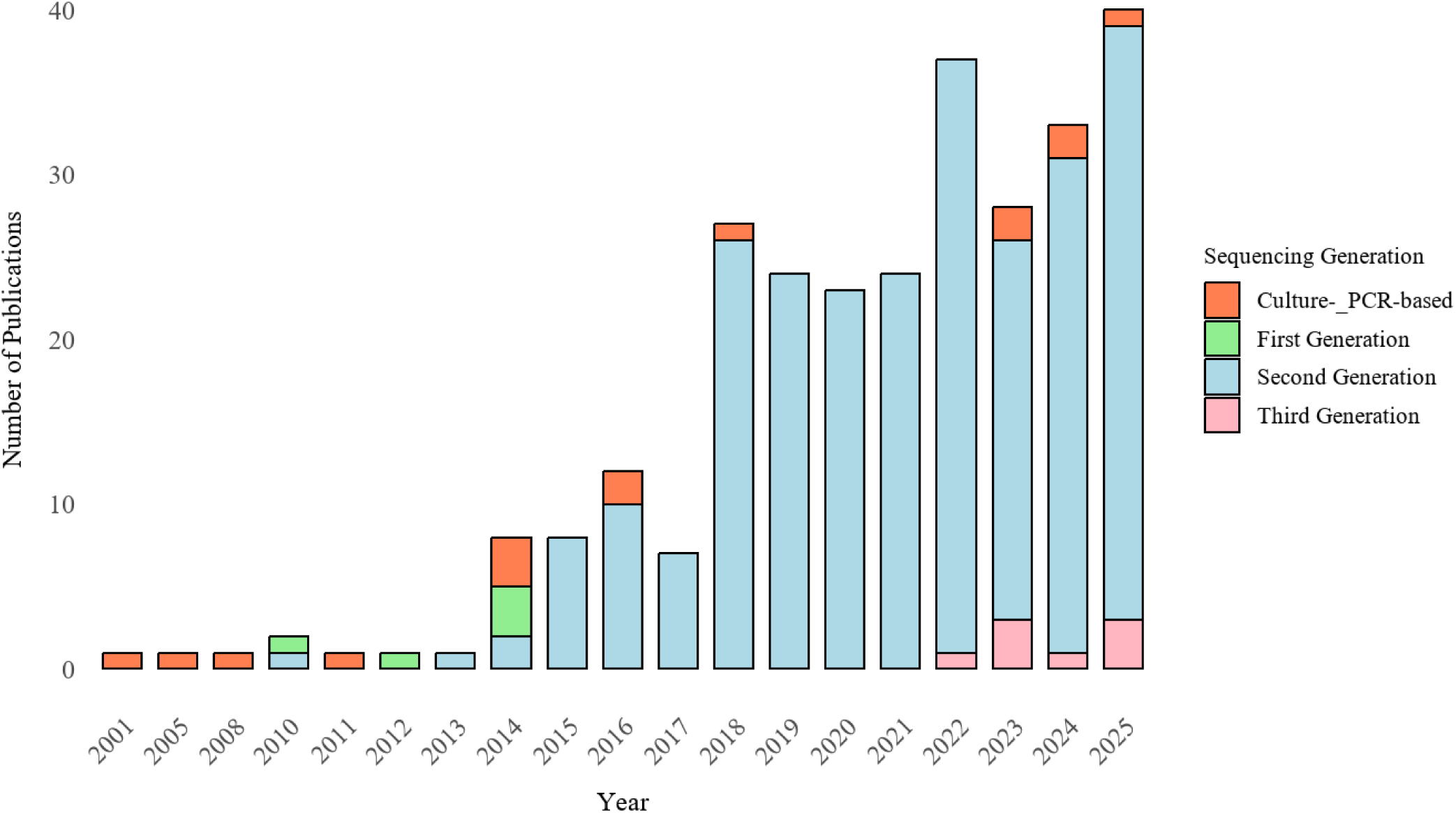
The number of scientific articles published per year from 2001-2025 (years with zero publications are not shown; 2002, 2003, 2004, 2006, 2007, 2009).

The four most common research scopes were Diet-related Impacts on Microbiota (100), Wild Environment and Habitat Influences on Microbiota (97), Microbiota Characterisation and Diversity (76), and Other Host-specific Factors (62) (Figure 3, S1) (Table SVI). Publication counts per scope (for years with at least one publication) ranged from 1 to 23, with a mean of 4.23, median of 3, standard deviation of 3.80, and interquartile range of 5. Importantly, these research scopes have directly contributed to primate conservation efforts (Table SVII).

**Figure 3:**
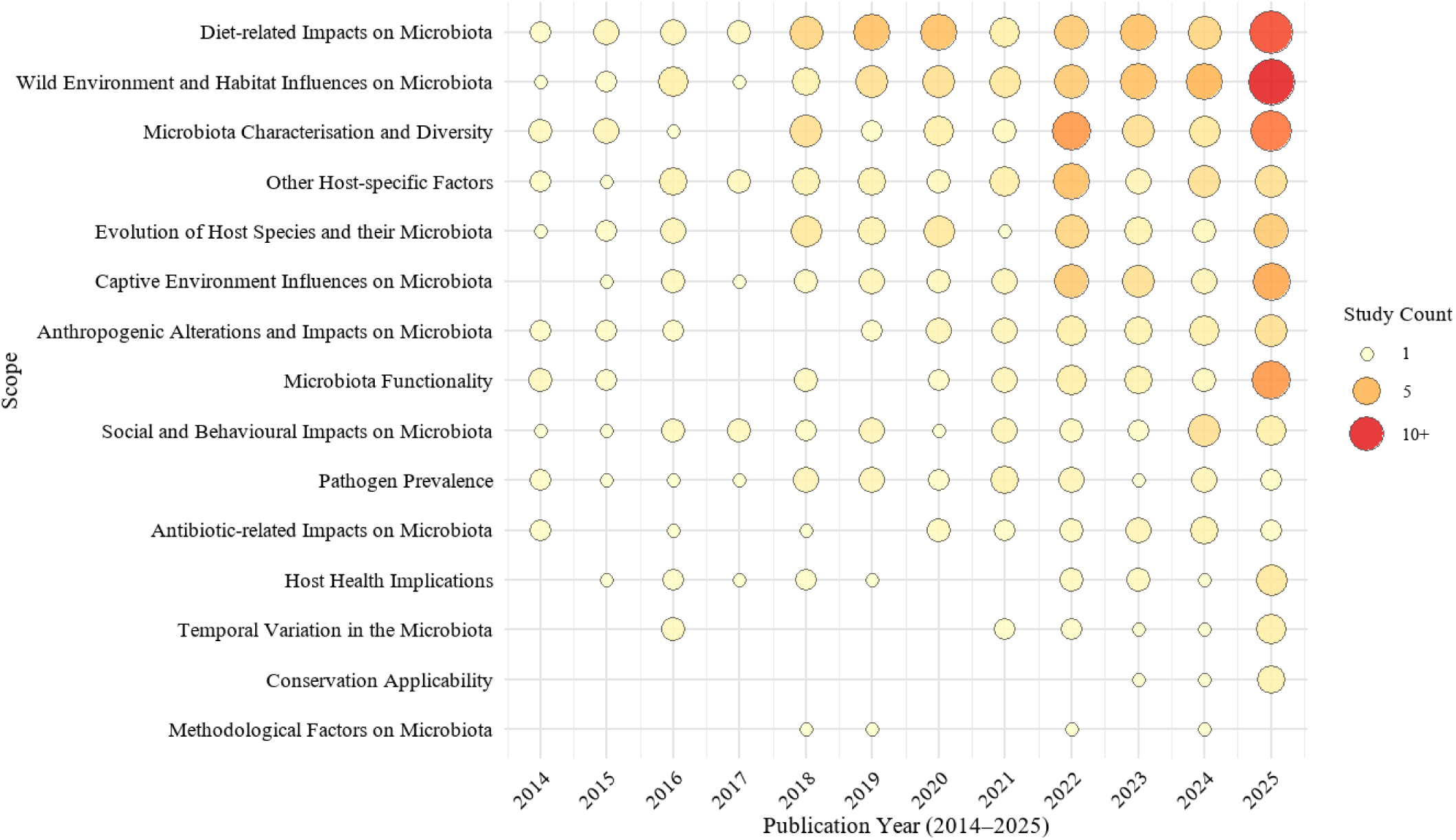
Bubble plot illustrates the frequency of each identified research scope in primate gut bacterial microbiome studies from 2014 onwards (full data is available in Figure SX). The size of each bubble represents the number of studies focusing on a specific scope each year, with larger bubbles indicating more frequent studies. The colour gradient, from yellow to red, reflects the increasing frequency of studies, with yellow representing fewer studies and red indicating higher study counts (Figure SX includes full dataset).

### Taxonomic coverage in primate gut bacterial microbiome studies

Coverage at the family level was high, with 15 of 16 (93.75%) primate families represented (Figure 4a); with Tarsiidae not covered. The mean number of studies per family was 25.1 (range 2-127), indicating broad but uneven coverage. The most studied families were Cercopithecidae (48.66%, n = 127), Hominidae (19.92%, n = 52), and Lemuridae (16.09%, n = 42) (Table SVIII). The proportion of species studied within each family was considerably uneven (Figure 5). One family has full species coverage (100% of species studied) (Daubentoniidae). Two families were extensively studied, with ∼76-86% of species sampled (Hominidae, Lemuridae). Two show moderate coverage (52-60%; Atelidae, Hylobatidae) and four with limited coverage (∼29-48%; Cercopithecidae, Indriidae, Callitrichidae, Lorisidae). Six families remain largely underrepresented, with ∼8-23% coverage (Cheirogaleidae, Galagidae, Cebidae, Aotidae, Lepilemuridae, Pitheciidae) and one has zero coverage (Tarsiidae), highlighting significant taxonomic gaps.

**Figure 4:**
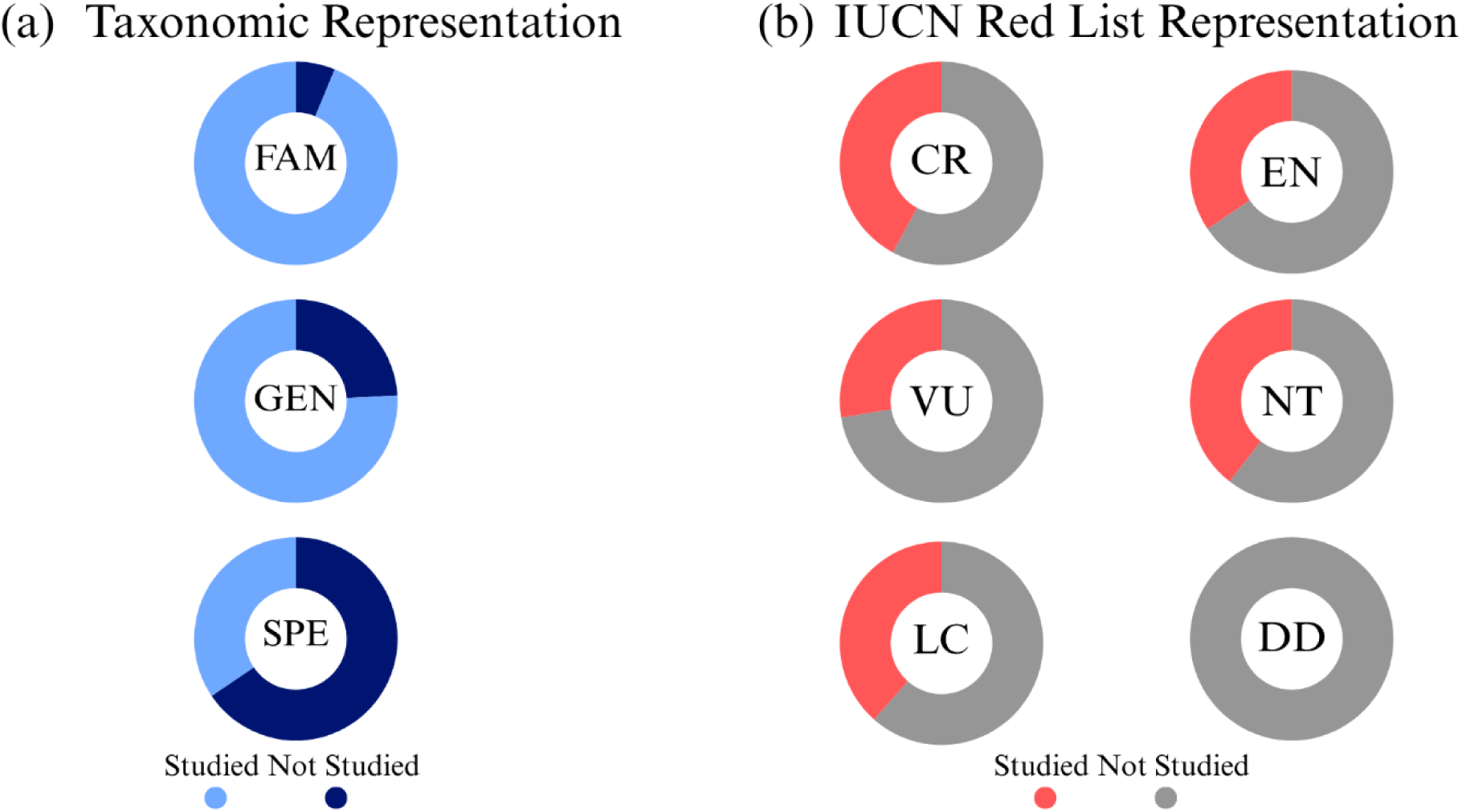
(a) The proportions of primates studied at the taxonomic levels of family, genus, and species; FAM = Family, GEN = Genus, SPE = Species. (b) The proportions of primate species studied from each IUCN Red List category; CR = Critically Endangered, EN = Endangered, VU = Vulnerable, NT = Near Threatened, LC = Least Concern, DD = Data Deficient. Figure created using Canva (https://www.canva.com/).

**Figure 5:**
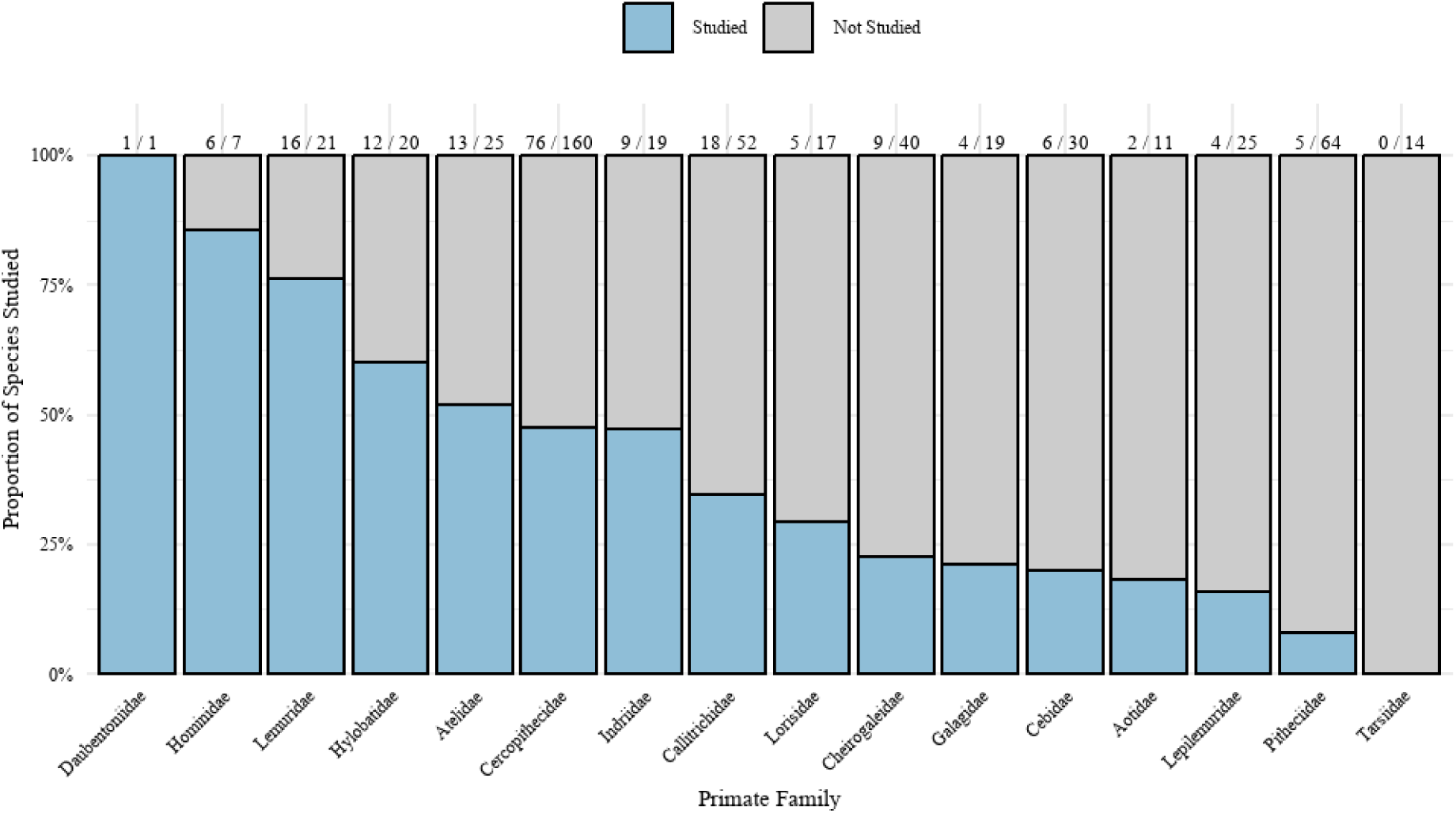
The proportion of species studied from within each of the recognised primate families.

Overall, 63 of 83 (75.90%) of primate genera were represented across studies (Figure 4a). The mean number of studies per genus was 9.52 (range 1-43), with seven of these genera represented by just a single study. The most studied genera were *Macaca* (16.48%, n = 43), *Rhinopithecus* (15.33%, n = 40), and *Gorilla* (13.41%, n = 35) (Table SIX). Regarding genera coverage (species studied per genus), 16 genera have full species coverage (100% of species studied) (*Papio*, *Mandrillus*, *Theropithecus*, *Procolobus*, *Nasalis*, *Callimico*, *Lagothrix*, *Varecia*, *Lemur*, *Prolemur*, *Symphalangus*, *Indri*, *Otolemur*, *Gorilla*, *Pan*, *Daubentonia*). Three genera were extensively sampled, with ∼75-83% coverage (*Eulemur, Rhinopithecus, Leontopithecus*). Moderate coverage was shown for 20 genera (∼50-72%) (*Ateles*, *Colobus*, *Callithrix*, *Galago, Oedipomidas*, *Hoolock*, *Propithecus*, *Pongo*, *Cercopithecus*, *Chlorocebus*, *Nomascus*, *Pygathrix, Semnopithecus*, *Miopithecus*, *Tamarinus*, *Cebuella*, *Mirza*, *Brachyteles*, *Nycticebus*, *Xanthonycticebus*). Limited coverage was shown for 11 genera (∼25-46%) (*Macaca*, *Cercocebus*, *Erythrocebus*, *Saguinus*, *Hylobates*, *Trachypithecus*, *Saimiri*, *Sapajus*, *Alouatta*, *Phaner, Hapalemur*). A further 13 genera remain largely underrepresented (∼6-24%) (*Cheirogaleus*, *Avahi*, *Leontocebus*, *Aotus*, *Piliocolobus*, *Microcebus*, *Presbytis*, *Cacajao*, *Cebus*, *Plecturocebus*, *Lepilemur*, *Mico*, *Pithecia*). Finally, 20 genera have zero coverage (*Allochrocebus*, *Lophocebus*, *Simias*, *Allenopithecus*, *Rungwecebus*, *Callicebus*, *Chiropotes*, *Cheracebus*, *Callibella*, *Allocebus*, *Paragalago*, *Galagoides*, *Sciurocheirus*, *Euoticus*, *Perodicticus*, *Arctocebus*, *Loris*, *Tarsius*, *Cephalopachus*, *Carlito*).

Of the 525 extant primate species recognised by the IUCN (2025a), just 181 species (34.48%) have been sampled to date (Figure 4a), representing substantial knowledge gaps. In addition, five hybrid taxa have been studied. The mean number of studies per species was 4.16 (range 1-33), with 67 of these species represented by just a single study. The most studied species were the Western lowland gorilla (*Gorilla gorilla*) (12.64%, n = 33), chimpanzee (*Pan troglodytes*) (12.26%, n = 32), ring-tailed lemur (*Lemur catta*) and the golden snub-nosed monkey (*Rhinopithecus roxellana*) (both 9.96%, n = 26) (Table SX).

### Geographic distribution of primate gut bacterial microbiome studies

Globally, samples were collected and studied across 425 distinct sites, including 286 wild sites, 135 captive sites, and four sites that were unclear (Figure 6). Most studies used samples collected in Asia (35.86%, n = 109), followed by Africa (33.55%, n = 102), North America (18.42%, n = 56), South America (7.57%, n = 23), Europe (3.95%, n = 12), Oceania (0.66%, n = 2), and one study did not state the country or region of sampling (F igure 7a). Overall, primates were sampled across 52 countries (Table SXI), with the top three being China (29.50%, n = 77), Madagascar (18.01%, n = 47), and the USA (13.03%, n = 34) (Figure 7b).

**Figure 6:**
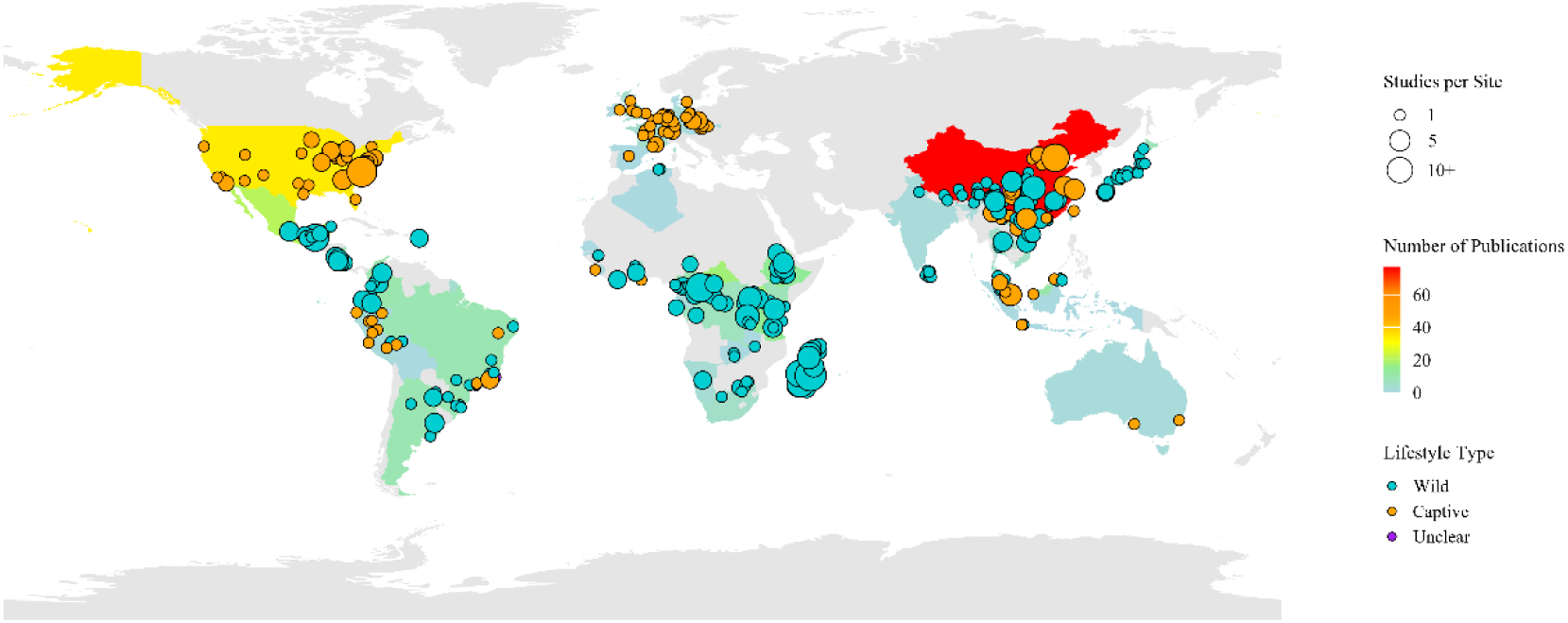
Global distribution of primate gut bacterial microbiome study sites. Country colour fill represents the total number of publications per country. The dot size represents the number of studies per site. The dot colour represents the lifestyle type of the sampled primates (i.e. wild or captive). Grey-shaded countries represent countries with no reported primate sampling locations in the reviewed literature.

**Figure 7:**
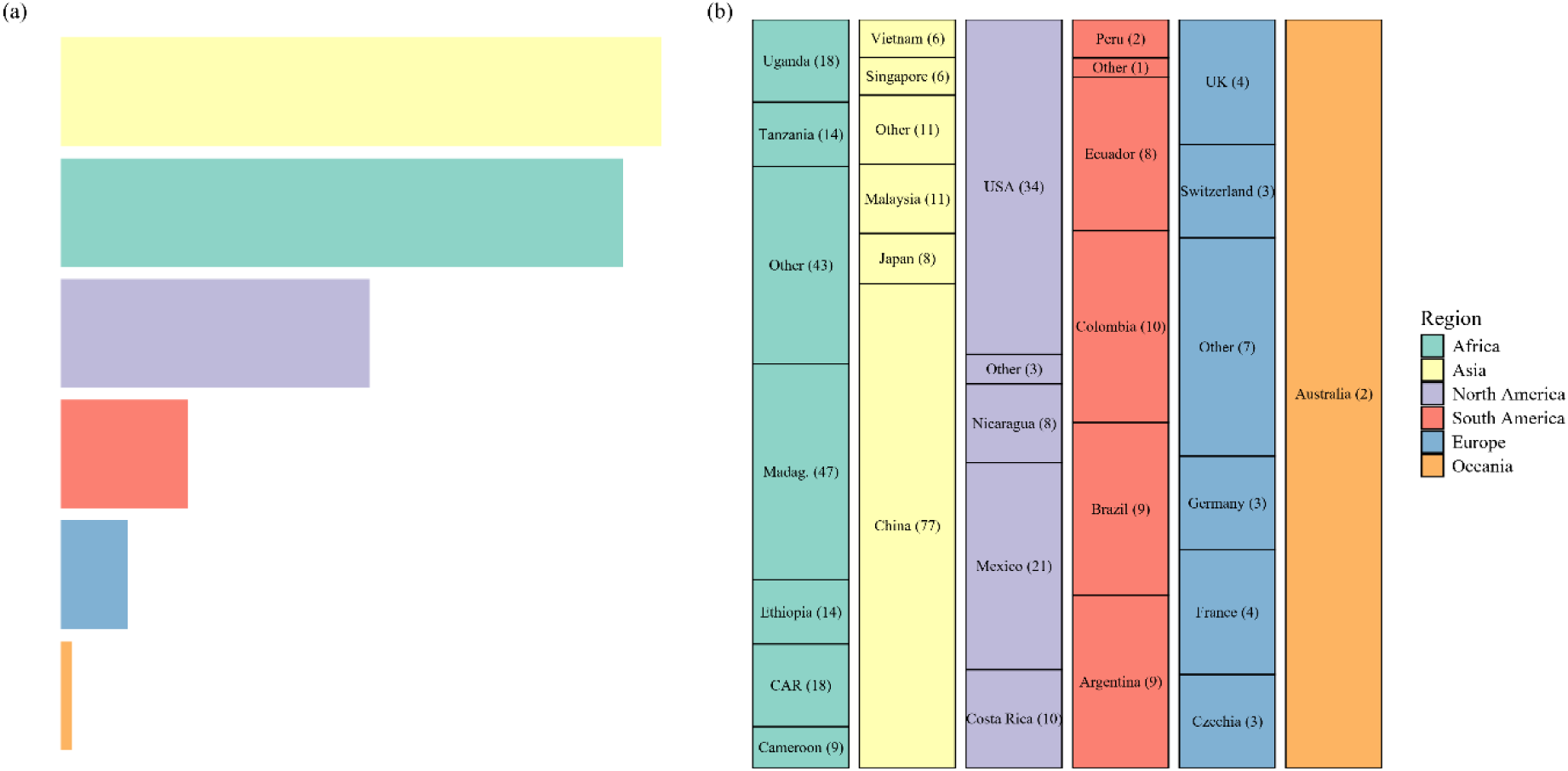
(a) Country representation within the six geographical regions in our dataset. Bars show the number of studies per country within each region, represented as a proportion of the region’s total. Within regions, countries contributing <5% were grouped as “Other”. A single study could include multiple distinct countries from the same region. (b) The proportion of all studies that each region was sampled in. Madag. = Madagascar; CAR = Central African Republic.

### Coverage across lifestyle contexts and IUCN Red List threat categories

Across studies, 83.14% (n = 217) included wild primates and 37.55% (n = 98) included captive primates. Of the 186 primates and hybrid taxa included in these studies, 67.20% have been sampled in the wild and 32.80% have only ever been sampled in captivity (Table I). Overall, just 23.81% of all IUCN-recognised species have been sampled in their natural wild habitat (Table I).

**Table I:** The count, and percentage, of all primate species that have been sampled across wild and captive lifestyle contexts.

| **SAMPLING HISTORY** | **SPECIES COUNT** | **PERCENTAGE (SPECIES AND HYBRIDS IN DATASET: 186)** | **PERCENTAGE (ALL RECOGNISED SPECIES ON THE IUCN: 525)** |
| --- | --- | --- | --- |
| Sampled in the wild | 125 | 67.20 | 23.81 |
| Sampled in captivity | 118 | 63.44 | 22.48 |
| Sampled in the wild AND captivity | 57 | 30.65 | 10.86 |
| Only sampled in the wild | 68 | 36.56 | 12.95 |
| Only sampled in captivity | 61 | 32.80 | 11.62 |

Threatened primates (Critically Endangered (CR), Endangered (EN), Vulnerable (VU), Near Threatened (NT)) were included within 83.91% of studies (n = 219). In isolation, CR species were included within 34.48% of studies (n = 90), EN species in 55.17% (n = 144), VU species in 22.61% (n = 59), NT species in 9.58% (n = 25), and Least Concern (LC) species in 31.03% (n = 81). Considering total species coverage within each threat category, 37 of 88 CR species (42.05%), 50 of 145 EN species (34.48%), 32 of 116 VU species (27.59%), 17 of 43 NT species (39.53%), 45 of 117 LC species (38.46%), and 0 of 14 Data Deficient (DD) species have been studied to date (Figure 4b), indicating similar coverage across most categories.

### GLMM and PGLMM: do primate traits influence species study effort?

Unknown population trend showed a significant positive effect on species study count in generalised linear mixed model one (GLMM1) (β = 0.69, SE = 0.32, z = 2.16, p = 0.03), whereas unknown trophic guild and unknown IUCN status showed significant negative effects on species study count in GLMM1 (unknown trophic guild: β = -2.20, SE = 0.64, z = -3.45, p = 0.00; unknown IUCN status: β = -5.35, SE = 1.24, z = -4.30, p = 0.00).

Body mass (kg) and region count (number of regions a species is distributed across; including native and introduced) showed significant positive effects on species study count in GLMM2 (Body mass: estimate (β) = 0.27, standard error (SE) = 0.13, z = 2.03, p = 0.04; region count: β = 0.21, SE = 0.09, z = 2.38, p = 0.02) and phylogenetic GLMM (PGLMM) (Body mass: β = 0.03, SE = 0.01, z = 3.04, p = 0.00). Conversely, the frugivore trophic guild showed a significant negative effect on species study count in GLMM2 (β = -0.78, SE = 0.39, z = -2.00, p = 0.05). See Table SXIV, SXV, SXVI, and Figures S2, S3, S4 for full data and further interpretation.

### Methodological variables reported across studies

Across reporting studies, the average number of samples analysed was 260.2 (79.31%, n = 207; range = 2-16,234, SD = 1461.2), and the average number of individual hosts sampled was 58.5 (61.69%, n = 161; range = 2 -585, SD = 85.9). Surprisingly, 20.69% of studies did not state the number of samples analysed, and 38.31% omitted the number of unique individuals sampled.

The most common sample preservation method prior to freeze-storage were cold storage methods (e.g., dry ice, liquid nitrogen, ice packs) (35.63%, n = 93), followed by stabilising buffers (e.g., RNALater, OMNIgene.GUT buffer) (31.42%, n = 82), then ethanol (70-100%) (21.84%, n = 57). Less common methods include using transport media, FTA cards, lysis buffers, drying methods, and fixative buffers (Table SXII).

Faecal samples were by far the most common sample type (93.49%, n = 244), followed by rectal samples (6.90%, n = 18), stomach samples (2.68%, n = 7), intestine samples (2.30%, n = 6), and then mixed samples (0.38%, n = 1) (some studies used more than one sample type).

The most common sequencing technology used across studies was Illumina (78.16%, n = 204), followed by 454 (Roche) (6.51%, n = 17), and then PacBio (2.30%, n = 6). Other sequencing technologies included Sanger, Ion Torrent (PGM), Oxford Nanopore Technologies (ONT), and MGI Tech. Additionally, in 14 studies (5.36%) sequencing technologies were not used; traditional PCR and/or culturing-based approaches were used instead (Table SXIII).

Among the Next Generation Sequencing (NGS) approaches employed across studies, amplicon sequencing was the most used (83.14%, n = 217), followed by shotgun metagenomics sequencing (21.07%, n = 55), metatranscriptomic sequencing (0.77%, n = 2), and shotgun pyrosequencing (0.38%, n = 1). In 16 studies (6.13%), an NGS approach was not utilised. Within the studies that used amplicon sequencing, the 16S gene was overwhelmingly the marker of choice (98.62%, n = 214), while the VT, cpn60, gyrB, invA, and ipaH markers were all used in one study each (0.46%). Some studies employed multiple sequencing technology, NGS approaches, or amplicon markers.

## DISCUSSION

### Drivers of uneven taxonomic coverage in primate gut bacterial microbiome research

A range of biological, ecological, logistical, and historical factors likely contribute to the uneven distribution of research effort across primate taxa, mirroring patterns seen in broader primatological research. Body mass and geographic range were consistent, positive predictors of species study count across models.

Larger-bodied species receive greater research effort, likely because their size, visibility, and behaviour facilitate field sampling. We found that genera such as *Gorilla*, *Mandrillus*, *Papio*, and *Pan* are extensively covered, whilst many small-bodied genera remain underrepresented (*Mico, Microcebus*), or only studied in captivity (*Cebuella*), or stilll unstudied (*Callibella*, *Carlito*). This finding is consistent with previous work demonstrating that larger-bodied primates are disproportionately studied (Junker et al., 2020; Ellison et al. 2021). Logistically, in addition to host-detectability, the size and quantity of their faeces may influence sampling success. Larger faeces are easier to track and locate, particularly when defecation occurs from a primate high in the canopy or when it falls within dense leaf litter (Burch, pers. comm.). Challenges may therefore compound when attempting to sample from small-bodied primates, both in tracking individuals until defecation and in locating smaller quantities of faecal material.

Geographic distribution (region count) also emerged as a significant positive predictor of species study count. This finding could be the result of increased accessibility and sampling opportunities rather than intrinsic biological importance of the species. If a species is distributed across multiple regions, it is more likely that multiple research groups could independently focus on populations of the same species across multiple locations. Furthermore, if a species is distributed across more than one region, it reduces the likelihood that the species will suffer from inaccessibility stemming from political instability or war.

Aside from these consistent positive predictors, other traits showed weaker, non-significant, effects. For example, models that included diel activity suggested a positive effect of diurnality and a negative effect of nocturnality on species study count. Taxonomic coverage broadly aligns with these trends too: five of the seven families with the least coverage are nocturnal primate families in the Aotidae, Cheirogaleidae, Galagidae, Lepilemuridae, and Tarsiidae. The remaining nocturnal families with greater coverage are Lorisidae and Daubentoniidae. Still, fewer than 30% of lorisid species have been sampled. As for Daubentoniidae, this family contains just a single species, the aye-aye (*Daubentonia madagascariensis*), which has only ever been studied in captivity. These patterns are consistent with previous findings that nocturnal or cryptid species are underrepresented (Junker et al., 2020). Practically, nocturnal primates are difficult to observe from the ground at night (Moore et al., 2021), and successful faecal sample collections in low-light conditions increases the difficulty. While these challenges can be mitigated by sampling nocturnal species in captivity, increasing intra-family/genera coverage, several studies have demonstrated that sampling nocturnal primates in the wild is feasible (e.g., Aivelo et al., 2016; Aivelo & Norberg, 2018; Greene et al., 2019; Perofsky et al., 2019; Wasimuddin et al., 2019), albeit logistically demanding and therefore less common.

Similarly, locomotion type (arboreal, terrestrial, or both) may influence detectability; however, it was not supported as a significant predictor in models where it was included. Arboreal primates may be more challenging to sample due to reduced visibility, and the primates often defecate from 10-30 metres above ground, with much of the faeces lost within tree branches and leaves (Orkin et al., 2016). Additionally, arboreal primates may travel via routes that are less accessible for researchers travelling on the forest floor. Additional factors such as social structure and group size may also influence detectability. Many of these traits are likely cumulative; species that are small-bodied, nocturnal, arboreal, and live in small groups may be particularly challenging to sample.

GLMM2 indicated a negative effect of the frugivorous trophic guild on species study count. Frugivorous primates, in comparison to folivores, are particularly challenging to study due to larger range sizes, lower population densities, rapid locomotion, and lesser accessibility associated with habitat suitability/tolerance to habitat alteration (Hawes & Peres, 2014). Hence, this pattern may reflect an indirect sampling bias related to other species traits, since it is unlikely that there is a direct bias against frugivore primates themselves.

Sampling accessibility via established research programs likely shapes patterns in taxonomic coverage. Hernandez et al. (2020) found that genomic studies often focus on species with extensive prior research histories. Similarly, many studies in our dataset targeted species repeatedly at a single site (e.g., *Alouatta pigra* at Palenque National Park, Mexico (Amato et al., 2013; 2014; 2015; 2016; 2017a; 2017b) or *L. catta* at the Duke Lemur Center, USA (McKenney et al., 2015; 2018; Greene et al., 2019; Bornbusch et al., 2022)). Accessibility may also take the form of access to previously collected biological samples or to existing sequence data housed in public repositories, both of which likely contributed to our findings since many studies in our dataset re-used existing sequence data (e.g., Pafčo et al., 2019; Amato et al., 2020; Mann et al., 2020; Sanders et al., 2023).

To further explore this effect, we included a measure of genus-level research legacy (WOS counts per genus, excluding studies included in this review) to assess whether historically well-studied genera are more likely to be targeted for gut bacterial microbiome research. Although GLMM2 indicated a positive trend, this effect was not statistically significant, suggesting limited support for research legacy as an independent predictor for study effort. Instead, our findings point toward the importance of established field sites and long-term study systems which enable repeated sampling and the expansion of ongoing ecological research into gut bacterial microbiome studies, often leading to the re-analysis of data generated from the same focal species over time.

GLMM1 identified significant effects associated with “unknown” trait categories (IUCN status, population trend, trophic guild). These results likely reflect artefacts of missing data, as this model retained all species by converting missing values to “Unknown”. However, they also highlight substantial gaps in ecological data across primates, which may limit the effectiveness of large-scale datasets for comparative analyses. Additional limitations include a substantial reduction in the number of species included in GLMM2 and PGLMM, since rows with missing data were excluded to reduced potential bias. This approach, however, may have introduced bias in the opposite direction by disproportionately removing poorly studied or data-deficient species.

Overall, a broad taxonomic coverage is seen at the family-level whereas at the finer scales, notable gaps are present across many taxonomic groups with limited intra-genus representation (i.e., several genera represented by a single species). Currently, 344 species (∼65.5%) are lacking information about their gut bacterial microbiome diversity, which could provide novel insights into wild primate population health and support maintenance of healthy captive populations.

### Studying familiar species in familiar contexts

Our findings revealed that whilst most studies include samples from wild primates, these studies have sampled relatively few species overall. Conversely, the lower proportion of captivity-based studies have focused on a relatively large proportion of the currently studied species. This skews our broader understanding of global primate gut bacterial microbiome diversity. Studies on the gut microbiome of captive primates can support primate conservation efforts in their own right, for example, these can improve husbandry practices, reduce diet-related health issues, and support the success of long-term captive breeding programs and can also guide pre-release management and support the long-term success of reintroduction programs (Frankel et al., 2019; Quiroga-González et al., 2021; Table SVII). Practicality-wise, captive individuals are more accessible, potentially improving sampling success of species that are more difficult to sample in the wild. However, captivity can substantially alter gut microbiome composition, diverging away from their wild conspecifics’ (Clayton et al., 2016). Furthermore, captive primate microbiomes may be correlated with cardiovascular disease and psychopathologies unlike those of their wild counterparts’ (Kummrow & Brüne, 2018; Dennis et al. 2019; O’Toole and Shiels, 2020). These microbial shifts are influenced by factors such as captive diet, environmental exposure, and husbandry practices, and may be especially pronounced in dietary specialists or species with naturally distinct microbial communities (Clayton et al., 2016; McKenzie et al., 2017; Asangba et al., 2019; Frankel et al., 2019). A strong argument can be made that primate taxa that have only ever been sampled in captivity still have poorly characterised gut bacterial microbiomes and cannot reliably be used as proxies for their wild conspecifics. Consequently, their relevance for conservation purposes remains limited.

Our results also highlighted that most studies included threatened species (CR, EN, VU, NT), and that species coverage per threat category was similar apart from DD primates. Similarly, Chen et al. (2023a) reported that most primate conservation publications targeted threatened species (∼68% of publications) but individual threatened species received uneven attention; with 0-3 publications for almost half of those species. Junker et al. (2020) likewise reported a similar pattern at the species-level, whereby conservation status did not strongly influence research effort in primate conservation intervention studies. On the contrary, Bezanson and McNamara (2019) reported that CR, DD, and No Evaluation (NE) species were included in just 18% of publications based on primate field research, revealing a lack of research effort on the most threatened, and knowledge devoid, species in the wild. The high study-level inclusion of threatened species in primate gut bacterial microbiome research indicates that primate gut bacterial microbiome studies are generally attentive to species of conservation concern. Attempts should be made to study the DD species to gain biological data which could then potentially be used to support future IUCN Red List category assessments.

### Geographic distribution of study sites: partial mismatch between primate species richness and research effort

Primates naturally occur in 90 countries, with two-thirds of all species concentrated in just four: Brazil (25.71%), the Democratic Republic of the Congo (DRC) (9.52%), Indonesia (12.38%), and Madagascar (20.19%); making these nations key priorities for primate conservation (Mittermeier, 1988; Estrada et al., 2017; IUCN, 2025b).

Global primate species richness is concentrated in a few hotspots: South America (the Amazon–Guianas and the Atlantic Forest), the Congo Basin and West/Central Africa, Madagascar, and much of Southeast and South Asia (e.g., Borneo, Sumatra, Indochina, and parts of India) (Figure 8) (Jenkins et al., 2013; Pimm et al., 2014; Biodiversitymapping, n.d.). When comparing global sampling effort to global primate species richness, a partial mismatch is seen. While Asia and Africa receive substantial attention that broadly matches their high richness, South America is markedly under-studied relative to its richness. North America and Europe are disproportionately represented despite lacking native species, highlighting a strong emphasis on captive-based studies (i.e., McKenney et al., 2014; McKenzie et al., 2017; Greene & McKenney, 2018; Li et al., 2020; Modrackova et al., 2021).

**Figure 8:**
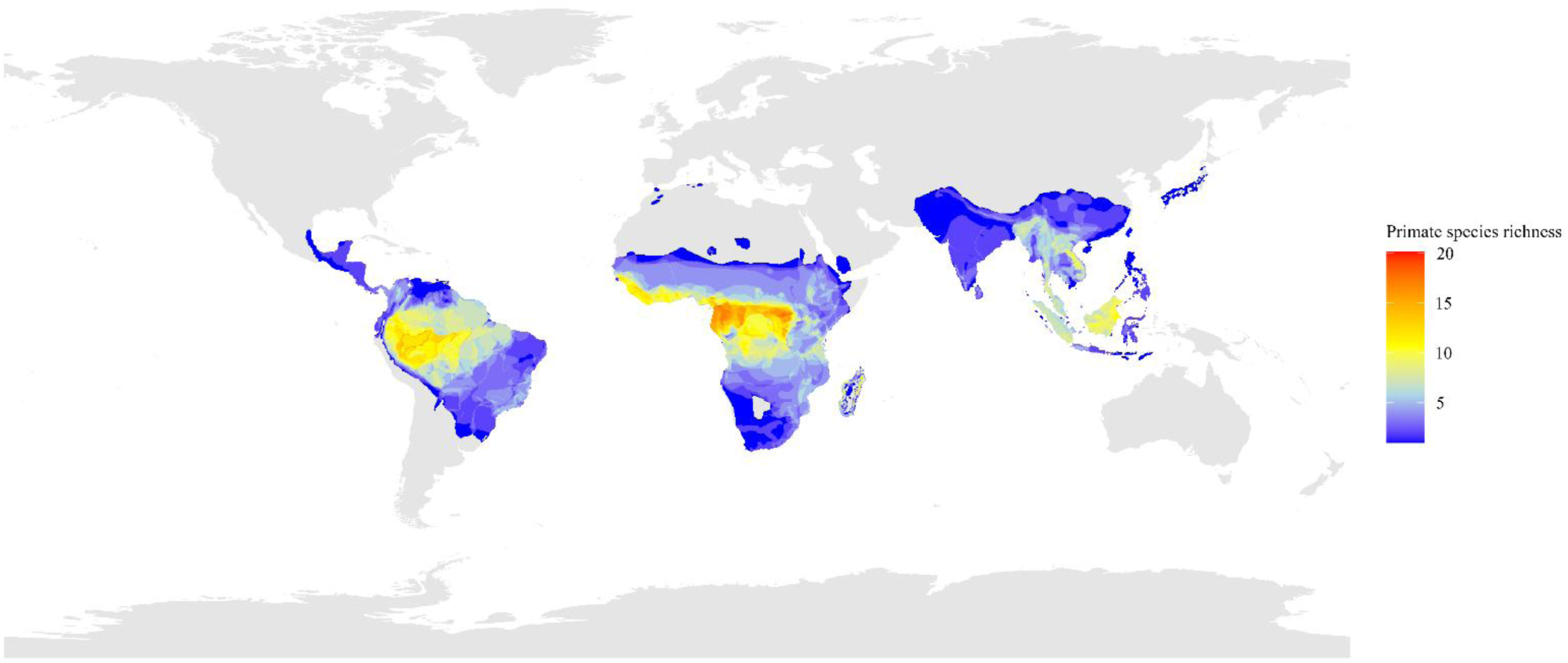
Global primate species richness. Data extracted from biodiversitymapping.org (Jenkins et al. 2013; Pimm et al. 2014; Biodiversitymapping, n.d.). Grey-shaded countries represent countries with no extant primate populations or where primate species richness was not resolved in the global dataset used.

Asia scored as the most sampled-within region, being led by China. This may be explained by the high species diversity coupled with the rapid expansion of primatology as a field of study in China (Garber, 2018). This expansion has been driven by a substantial increase in the number of individuals receiving advanced degrees (Fan & Ma, 2018; Garber, 2018). In 2017, this growth culminated in the establishment of the China Primatological Society (Garber, 2018). This growth likely accounts for the increasing publication output trend, with 94.81% of studies conducted in China published from 2018 onwards. In contrast, Indonesia is notably undersampled despite being a global hotspot for primate diversity with high conservation priority (Mittermeier, 1988; Grow et al., 2010; Estrada et al., 2017). This contrast is surprising considering that Indonesia was the second most common location for primate conservation-related publications between 1994 and 2019 (de Figueiredo Machado et al., 2023). Furthermore, a review that looked at studies focusing on zoonotic pathogens (protozoa, GI parasites, viruses, bacteria, fungi, and blood-borne parasites) conducted between 1965 and 2023 on primates in Asia found that Indonesia was the joint top with Thailand (25 studies each) (Patouillat et al., 2024).

The research attention given to Africa mirrors patterns in broader primate field research, whereby 55.5% of published primate field research was conducted across Africa (including Madagascar) (Bezanson & McNamara, 2019). Of the African nations, Madagascar was most featured in primate gut bacterial microbiome studies. This research effort is warranted, given both the high primate species richness and the unique primate assemblage found on this island. Madagascan primates are split across five families, most are endemic (with two species also present on the Comoro Islands), and all face severe extinction threats. These same reasons may explain why primate-conservation-related publications were most conducted across Madagascar between 1994-2019 (de Figueiredo Machado et al., 2023). Even in broader conservation-related research, Madagascar ranked fifth in the number of conservation science papers published between 1987-2017; behind South Africa, Kenya, Ethiopia, and Tanzania (Pototsky & Cresswell, 2021). In contrast, research efforts toward primates within the DRC has been minimal. Like Madagascar, the DRC can be considered a primate conservation priority country (Estrada et al., 2017). Yet, few studies sampled within the DRC and those that did focused solely on bonobos (*Pan paniscus*), leaving all but one taxon unstudied. Ecological research efforts within countries of the eastern Congo Basin, such as the DRC, face logistical issues, armed conflict, and political instability (van der Hoek et al., 2021), which may partly explain this research void.

South America was represented in only a small fraction of global studies (∼7.5%), which is surprising considering the high primate species richness throughout much of the region. The most contrasting comparison between primate species richness and research effort was seen in Brazil (especially in the Amazon region), which is one of the four high-priority areas for primate conservation (Mittermeier, 1988; Estrada et al., 2017). Unexpectedly, no publications to date have reported on the gut bacterial microbiome of primates inhabiting the Brazilian Amazon. Salvarani et al. (2025) reported that research into bacterial diseases in wild animals of the Amazon is still in its infancy, consequentially, there is a lack of knowledge of disease prevalence, transmission dynamics, and ecological consequences within this biome. This disparity is not restricted to biological sampling efforts, since ecological research in the Brazilian Amazon is primarily driven by logistics and human influence (Carvalho et al., 2023). Interestingly, the authors note that research probability declined on indigenous lands, which represents about 23% of the Brazilian Amazon. Prioritising field sampling in Brazil and adjacent Amazonian countries would address one of the clearest global mismatches between primate diversity and research effort.

Finally, studies across North America, Europe, and Oceania were driven by captive sampling. While these studies enable a more controlled environment for comparisons and longitudinal analysis, its application into species conservation may remain limited as previously mentioned. Targeted paired designs, i.e., captive–wild comparisons for the same species (e.g., Villers et al., 2008; Fogel et al., 2015; Torfs et al., 2025) could substantially increase conservation applicability.

Overall, research effort is biased, leaving megadiverse countries largely underrepresented. Most studies are concentrated in China, Madagascar, the USA, and Mexico. Whilst Madagascar is both species rich and well-studied, other key species rich countries like Brazil, Indonesia, and the DRC remain understudied. Addressing these geographic gaps with targeted sampling of unstudied species in understudied countries would broaden our understanding of global primate gut bacterial biodiversity. Currently, our understanding of primate gut bacterial microbiomes derives from data acquired from a small proportion of species. Moreover, one-third of these species’ microbiomes are altered by captivity. Expanding research efforts into unstudied wild primates is crucial for understanding microbial diversity and to inform *in-situ* conservation efforts. Ultimately, this will culminate with a more comprehensive view of the primate gut bacterial microbiome diversity across the full spectrum of primate species. Further in-depth discussion is included in the Supplementary Material.

### Temporal trends in primate gut bacterial microbiome research

Prior to 2010, just three studies had been published on the primate gut bacterial microbiome. Research scopes were narrow, due to low publication volume, with the most common research scope being Pathogen Prevalence (Figure S1). These studies focused on single-species identification (on more than three species of bacteria) and relied on culture-based and PCR assays e.g., in just two primate species–Eastern gorillas (*Gorilla beringei*) (Nizeyi et al., 2001; Kalema-Zikusoka et al., 2005) and ring-tailed lemurs (*L. catta*) (Villers et al., 2008). Both Nizeyi et al. and Kalema-Zikusoka et al. aimed to identify *Campylobacter*, *Salmonella*, and *Shigella* specifically, whilst Villers et al. aimed to identify five specific bacterial groups. Although informative for basic host–microbe interactions, the single-species identification approach systematically underrepresents unculturable taxa and limits inferences about community structure and diversity (Arnold et al., 2016).

Between 2010 and 2013, only another five studies had been published, focusing on seven different primate species: the ashy red colobus (*Piliocolobus tephrosceles*), red-tailed monkey (*Cercopithecus ascanius*), guereza (*Colobus guereza*) (Yildirim et al., 2010), the pygmy slow loris (*Xanthonycticebus pygmaeus*) (Xu et al., 2010), the indri (*Indri indri*) (Junge et al., 2011), the Western gorilla (*G. gorilla*) (Vlčková et al., 2012), and the black howler monkey (*A. pigra*) (Amato et al., 2013). In 2010, Sanger sequencing (Sanger et al., 1977) was utilised for the first time for the primate gut bacterial microbiome (Xu et al., 2010). Sanger sequencing (chain termination) is based on DNA template amplification and gel electrophoresis (Eren et al., 2022). The accuracy, robustness, and ease of use made it the most common DNA sequencing technology for years (Heather & Chain, 2016). The application of Sanger sequencing for primate gut bacterial microbiome research did not last long due to the rise of more cost-and-time effective sequencing technologies. During this period, 16S rRNA amplicon sequencing (hereafter, 16S sequencing) was also employed for the first time. While 16S sequencing is typically considered an NGS approach, early applications used both first-generation Sanger sequencers (Xu et al., 2010) and second-generation sequencers such as 454 (Roche) (hereafter, 454) (Yildirim et al., 2010). The 16S gene, abundant in bacteria and containing both conserved and hypervariable regions, has since become the most widely used molecular marker for microbial identification (Yu et al., 2019; Varliero et al., 2023). These qualities make it ideal for gut bacterial microbiome community characterisation across host species. By using primers that bind to conserved regions of the 16S gene to amplify intervening hypervariable regions, and sequencing amplicons, reads can be aligned to a taxonomic reference database, enabling much broader identification of microbial taxa than was possible with earlier culture-based or low-throughput PCR approaches such as polymerase chain reaction-denaturing gradient gel electrophoresis (PCR-DGGE) or quantitative polymerase chain reaction (qPCR). However, the accuracy of taxonomic assignments still largely relies on the robustness of the chosen database.

Following the early application of second-generation sequencing by Yildirim et al. (2010), there was a lag before its broader adoption; from around 2013 these technologies gained traction and rapidly became the dominant approach. The uptake of second-generation high-throughput sequencing from 2013 onwards removed much of the time and cost associated with using first-generation (or earlier) sequencing approaches, owing to massively parallel sequencing and substantially reduced per-base sequencing costs (Shendure & Ji, 2008; Metzker, 2010). This shift marked a major transitory point for the field, enabling more comprehensive community profiling and driving the subsequent rise in both the scale and scope of primate gut bacterial microbiome studies, with the first diet-related scope appearing in 2013 (Amato et al., 2013). However, the full impact of this transition became evident in the years that followed.

In 2014, a noticeable rise in publication frequency occurred with eight publications (Figure 2). This pattern signalled a shift away from traditional, low-throughput methods and contributed to the sustained growth of primate gut bacterial microbiome research. Research scopes also broadened beyond early focus on scopes such as pathogen prevalence and microbiota characterisation, encompassing studies that focused on scopes such as microbiota functionality, social and behavioural impacts on host microbiota, and antibiotic-related impacts on host microbiota (Figure 3, S1). However, at this stage studies were still somewhat reliant on first-generation sequencers and PCR- or culture-based approaches. Of these studies, 62.5% conducted 16S sequencing using first- or second-generation sequencers; the other 37.5% still utilised traditional culture based or PCR-assay methods (Figure 2). Also in 2014, a substantial increase in the number of primate species studied occurred. Fifteen species were studied–more than had been studied over each of the previous years since 2001 combined (just nine). This period also aligned with broader developments in primate molecular research, as 2014 marked a substantial increase in the availability of high-quality primate reference genomes and re-sequencing data from multiple individuals (Roos et al., 2025), which may have indirectly facilitated the methodological changes emerging in primate microbiome research. Davenport et al. (2017) also reported that in the decade prior, improvements in technology for collecting and analysing DNA sequence data made possible answering questions related to the human microbiome composition within and across body sites, and how these relate to other species’ microbiomes.

The broader uptake of second-generation sequencing technologies occurred in 2015. During this year, another eight articles were published but all were conducted using second-generation sequencers; split 50:50 by Illumina and 454. Illumina was first utilised by Fogel (2015), McKenney et al. (2015), Moeller et al. (2015), and Tung et al. (2015). Unlike Sanger sequencers, which sequence one DNA fragment at a time, Illumina sequencers (and 454) allow for simultaneous sequencing of millions of short DNA fragments (Hu et al., 2021); significantly increasing the speed, depth, and resolution of microbiome analyses. Illumina would go on to become the dominant sequencing technology in primate gut bacterial microbiome research (Supplementary File 1), whereas 454 would not be utilised for much longer due to the discontinuation of 454 (Bleidorn, 2016).

In 2015, Shotgun sequencing approaches were applied for the first time in primate gut bacterial research, offering greater taxonomic and functional resolution compared to 16S sequencing (Jovel et al., 2016). Unlike 16S sequencing, which can be biased by primer choice when targeting different hypervariable regions (Manus et al., 2024), shotgun metagenomics sequences all DNA in a sample, capturing bacteria, viruses, fungi, algae, (amongst other microbial taxa) and host DNA, enabling analyses into microbial community dynamics and functionality (Sharpton, 2014). Early studies included Tung et al. (2015) on yellow baboons (*Papio cynocephalus*) and Xu et al. (2015) on black snub-nosed monkeys (*Rhinopithecus bieti*) (shotgun pyrosequencing). Shotgun pyrosequencing, essentially the same as shotgun metagenomics sequencing but conducted on 454 sequencers, is now obsolete following the discontinuation of 454 (Bleidorn, 2016). Functional assessments using metagenomics, metabolomics, and metatranscriptomics alongside 16S are crucial for understanding microbial impacts on primate health (Clayton et al., 2018). Despite these advantages, 16S sequencing remains the most common NGS approach in primate gut bacterial microbiome studies to date (Supplementary File 1).

More recently, third-generation sequencers have been utilised for primate gut bacterial microbiome research (Figure 2). The first study to use a third-generation sequencer was published in 2022 by Ying et al. Unlike second-generation sequencers (e.g., the 454 (Roche) GS FLX Titanium or the Illumina MiSeq series sequencers), which rely on short amplicons and whose choice can influence results, third-generation sequencers (e.g., the ONT MinION sequencer) allow sequencing of full length 16S rRNA genes and can reduce reliance on PCR amplification, although small or targeted amplification steps are still possible. By reducing or eliminating PCR amplification during library preparation, third-generation sequencing technologies allow single-molecule sequencing (Bleidorn, 2016) and can achieve a finer taxonomic resolution than approaches targeting shorter hypervariable regions (Myer et al., 2016). Ying et al. (2022) used a PacBio third-generation sequencer, followed by Li et al. (2023), Wang et al. (2023), Long et al. (2024), Xia et al. (2025), and Zhou et al. (2025). ONT is another of the prominent third-generation sequencing companies but has so far only been used twice for shotgun metagenomics sequencing (Sanders et al., 2023; Schweikhard et al., 2025). As these technologies become more cost-effective and accessible, third-generation sequencing will likely become increasingly more prominent in primate gut bacterial microbiome research.

The shift from culture-based methodological approaches to high-throughput third-generation sequencing parallels the rapid expansion of mammal microbiome research more broadly. On Google Scholar, primate-based studies on the microbiome accounted for approximately half of the total mammal microbiome literature in the 10 years prior to Amato and Stumpf’s (2019) review, which included an increase from 236 to 913 articles published per year. This growth represents the expanding toolkit, a more widespread understanding of the potential of these microbiome insights, the greater availability of technologies, and the continued interest in primates as study subjects. Yet, despite the general increasing trend in publication frequency (which peaked in 2025 (38 articles; Figure 2)), several methodological inconsistencies remain.

### Standardisation: enhancing the applicability of gut bacterial microbiome research to support primate conservation efforts

Our results indicate variations in sampling effort, preservation methods, and reporting standards, which hinder comparability and limit the ability to draw robust conclusions at broader ecological scales. Fundamental aspects of study design, such as determining the appropriate number of samples to collect or how many distinct individuals to sample, are difficult to ascertain through literature review alone.

Given the considerable variation in sample sizes and numbers of individuals sampled across studies, it remains challenging to recommend a definitive “correct” sample size. Since potential sources of bias are lower for samples sizes that are five or greater, a minimum of five samples per host species has been recommended for comparative gut microbiome studies (Degregori et al., 2025). We suggest that researchers maximise sample sizes within the constraints of available resources to enhance the depth and robustness of their data to optimally address their specific research scope(s), whilst minimising potential biases (Degregori et al., 2025), and to always report the number of samples analysed in publications. The number of individual hosts to sample depends on study design. Longitudinal studies may require fewer hosts when repeated sampling is conducted over time, whereas shorter-term studies may benefit from sampling more hosts to capture population-wide microbiome profiles. Critically, explicit and consistent reporting of these variables is essential for assessing data robustness and enabling reliable cross-study comparisons.

During the data extraction process for this review, certain data were less accessible than expected; particularly data relevant to support primate conservation. For example, one publication, reported only the genus of the sampled species (for seven genera in total), omitting the full binomial nomenclature. The lack of taxonomic specificity limits the use of the study for species-specific conservation efforts as it constrains other researchers in their interpretation of the data.

We found inconsistencies in preservation methods across studies prior to freeze-storage. The gold standard is immediate freezing at -80C to maintain DNA quality and minimise microbial bias (Nearing et al., 2021), but this is often impractical in remote field sites. In such cases, stabilising methods can maintain samples until freezing is possible. For example, Song et al. (2016) reported that 95% ethanol, FTA cards, and the OMNIgene.GUT kit preserved samples at ambient temperatures for up to eight weeks, with microbial changes within normal technical variation. Our findings show most studies used stabilising methods, cold storage methods, or ethanol, and some combined approaches. Using a single preservation method per study is recommended to avoid batch effects (Song et al., 2016). We further recommend consulting comparative studies on faecal preservation methods in mammals (e.g., Hale et al., 2015; Song et al., 2016; Chen et al., 2019; Ma et al., 2020; Burnham et al., 2023), and reporting preservation methods explicitly to improve comparability and interpretation of results.

Conversely, some aspects of methodological approaches have been very consistent such as sample type, sequencing technology, and NGS approach. Most studies analysed faecal samples, a pattern reported previously by Clayton et al. (2018). The main advantage of using faecal samples is that they can be collected non-invasively, causing minimal disturbance to the primates. In contrast, rectal samples require some physical restraint, whilst stomach and intestinal samples generally require post-mortem collection. This consistency is important for cross-study comparison, since microbial profiles can vary substantially depending on the sample type (Videvall et al., 2018; Pietroni et al., 2025). However, faecal samples may not accurately represent the gut microbiome of the entire gastrointestinal tract (Yasuda et al., 2015; Clayton et al., 2018), which may be something to consider depending on research scope.

Most studies employed Illumina sequencers and 16S sequencing, the use of which can provide important baseline data that characterise the gut bacterial microbiome of each primate species. Overtime, we may see a shift with ONT and PacBio becoming more prominent with third-generation sequencers becoming more cost-effective. The use of shotgun metagenomics and metatranscriptomics will likely play an increasingly important role in this field of research, expanding study scopes to explore the functional roles that bacterial communities provide for their hosts. At a time when primate populations are declining globally due to increasing anthropogenic pressures, understanding how host-associated microbiomes function and respond to environmental change can provide valuable insights into primate resilience and conservation.

To support the conservation applicability of primate gut bacterial microbiome research, we propose the standardised reporting of a minimum set of data in primate microbiome studies (Table II). Ideally, these metadata should be reported in the main text of the article, rather than confined to the supplementary material or external citations. This would enhance transparency, facilitate reproducibility, and push towards advancing microbiome research in primatology by promoting greater methodological consistency.

**Table II:**
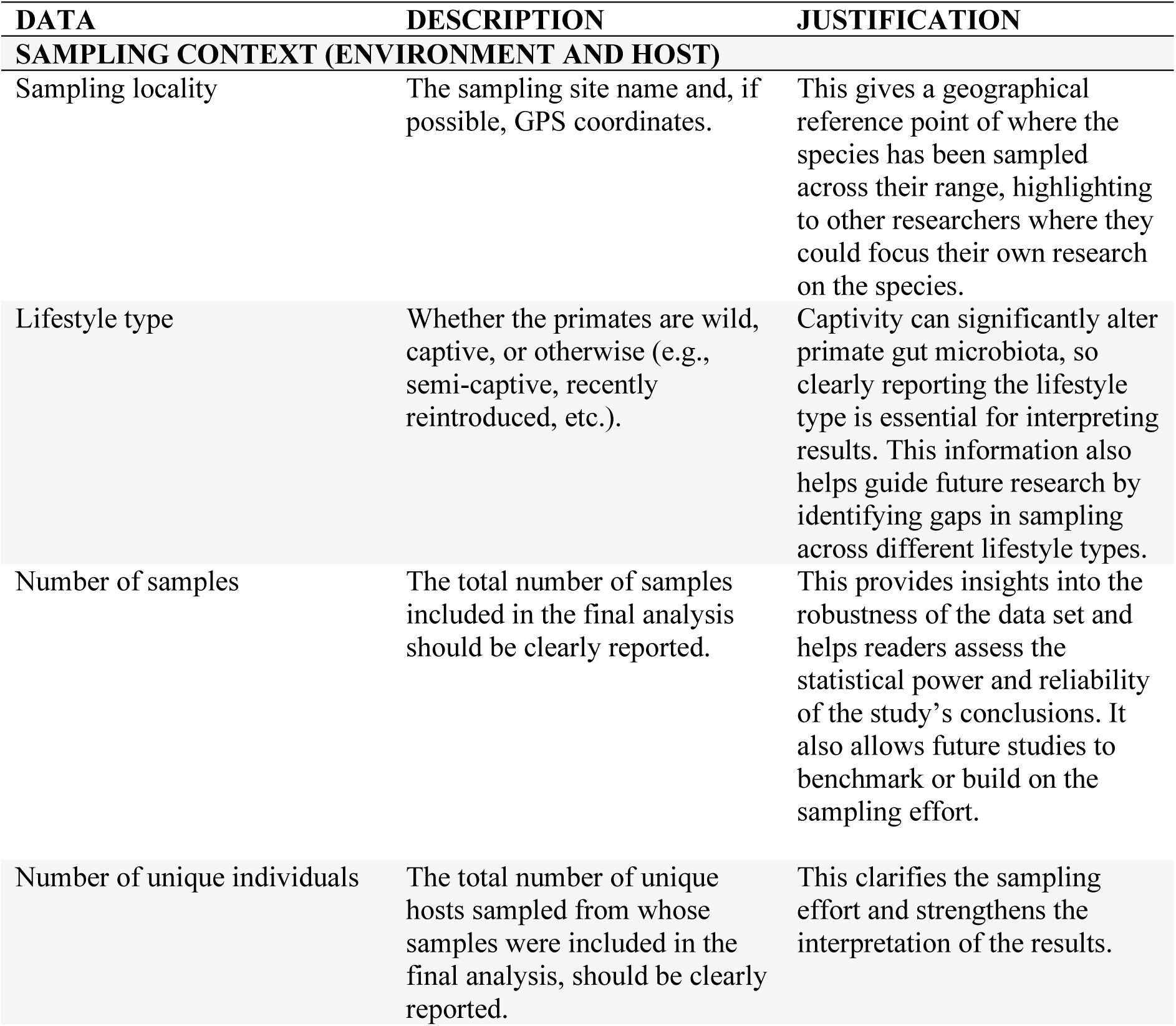
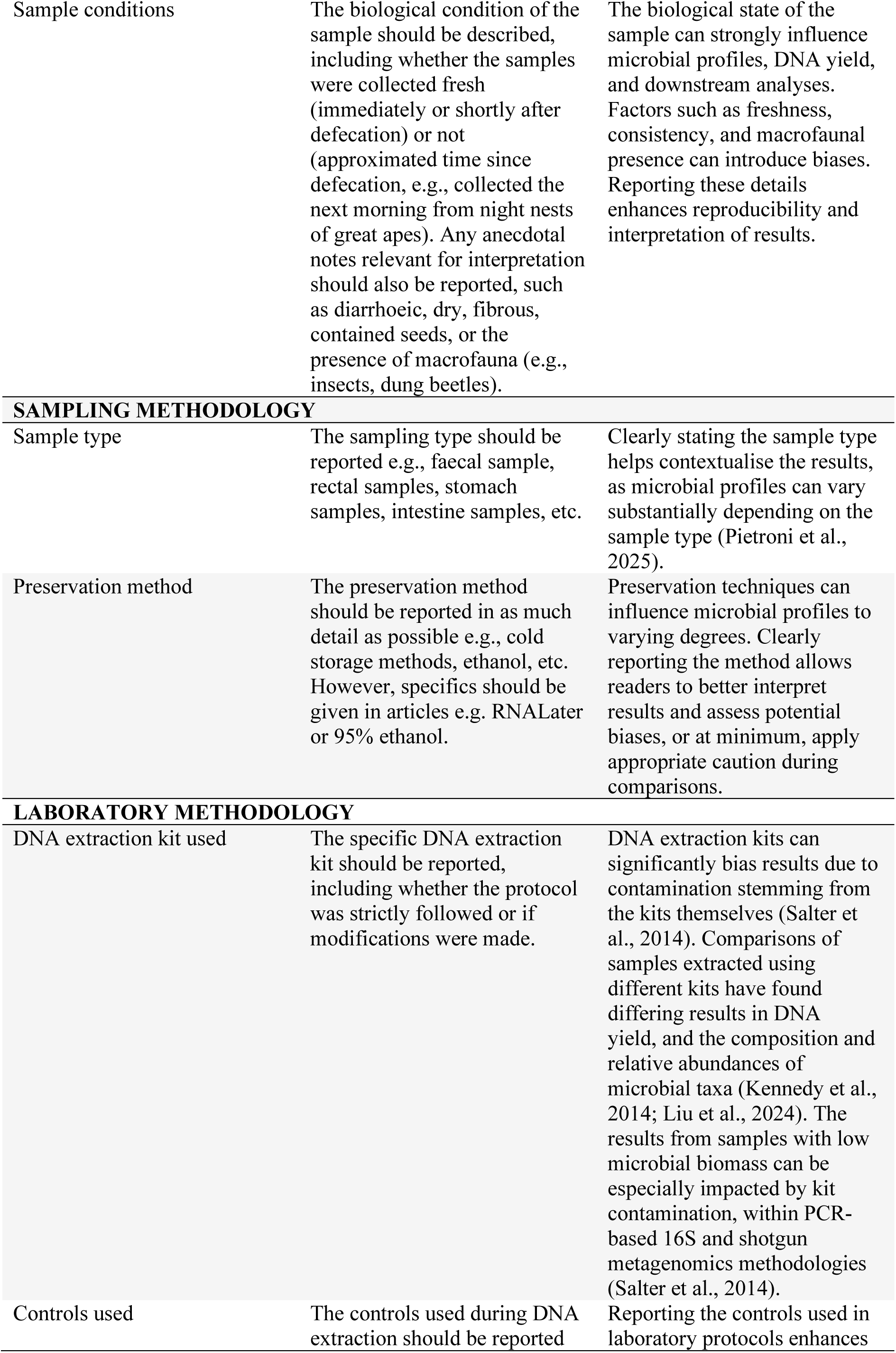
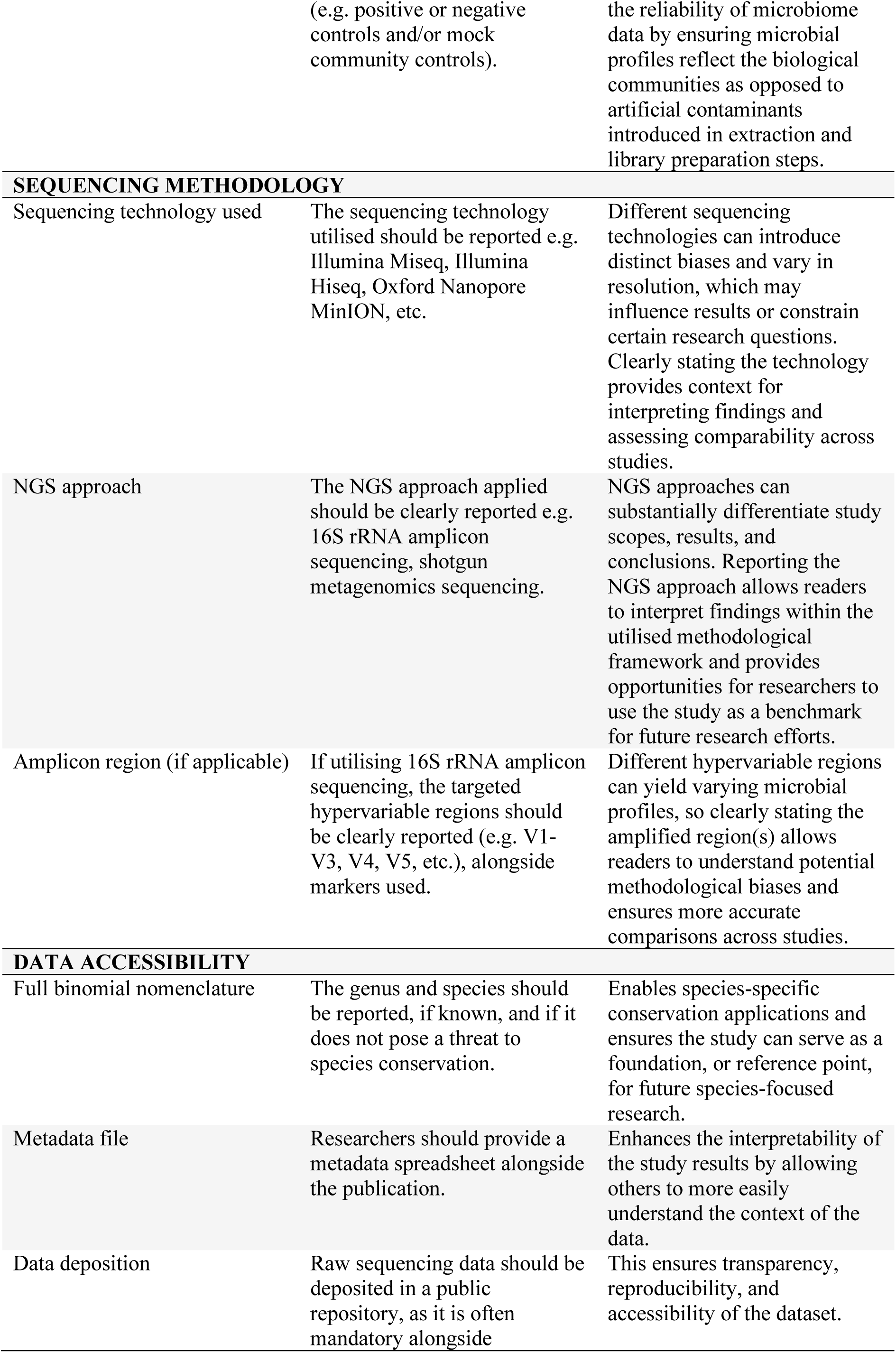
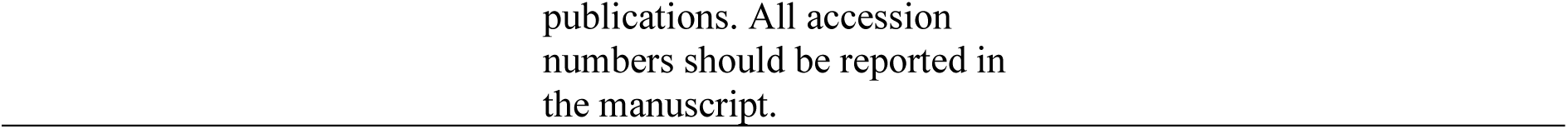
The proposed minimum set of data to be reported in future primate gut microbiome studies.

## CONCLUSION

The path towards a more comprehensive understanding of the global primate gut bacterial microbiome diversity lies in addressing the taxonomic gaps at the species level, particularly among those with low intra-family representation (i.e., Cheirogaleidae, Cebidae, Aotidae, Galagidae, Lepilemuridae, Pitheciidae, Tarsiidae). Research efforts should likewise address geographic gaps, focusing on the more threatened and DD primates within the megadiverse key-priority areas such as Brazil, the DRC, Indonesia, and Madagascar. Importantly, this review proposes the standardised reporting of a minimum set of data for future primate gut microbiome studies. Transparency is essential for standardisation efforts because it facilitates reproducibility, enhances cross-study comparisons, and makes information more accessible, promoting the gradual convergence of research methodologies over time. Collectively, these steps are essential for ensuring that primate microbiome research can meaningfully inform conservation strategies.

## Data Availability Statement

R scripts and Supplementary Files 1-5 will be made publicly available upon acceptance of this manuscript for peer-reviewed publication.

## Acknowledgements

We thank our collaborators from the Federal University of Mato Grosso (Sinop), the NGO Instituto Ecótono, the NGO Muriqui Project of Caratinga, and the Federal University of Rio de Janeiro for their collaboration on related microbiome research projects and discussions that informed this review. We also acknowledge the University of Salford Doctoral School Fund for supporting the fieldwork associated with related microbiome research that informed aspects of this review. Finally, we thank Dr Laura Brettell and Dr Marina Duarte for their thoughtful review of an earlier version of this manuscript, whose comments substantially improved it.

