## Supplementary Materials for "The primate gut bacterial microbiome: a systematic review of research methodologies, taxonomic coverage, and conservation implications"

### METHODS

#### Methods: tables

Table SI: Keywords and Boolean operators used to retrieve scientific articles on Web of Science and Scopus. For both databases, the Boolean operator AND was used between the primate, microbiome, and conservation keyword blocks.

| DATABASE | FIELD CODE | KEYWORDS FOR PRIMATES | KEYWORDS FOR MICROBIOME | KEYWORDS FOR CONSERVATION |
|---|---|---|---|---|
| Web of Science | All Fields | (“primate*” OR “monkey*” OR “ape*” OR “lemur*” OR “loris*” OR “tarsier*”) | (“microbio*” OR “microbe*” OR “bacterial communit*” OR “microflora”) | (“conservation”) |
| Scopus | Title-Abs-Key | ( primate* OR monkey* OR ape* OR lemur* OR loris* OR tarsier* ) | ( microbio* OR microbe* OR bacterial communit* OR microflora ) | ( conservation ) |

Table SII: Inclusion and exclusion criterion with justifications.

| CRITERION | INCLUSION | EXCLUSION | JUSTIFICATION |
|---|---|---|---|
| 1. Publication type | Peer-reviewed journal articles | Dissertations and theses | Avoid duplication of work already published in journals (Astiazarán-Azcárraga et al., 2024). |
| 2. Data type | Studies presenting primary data (newly generated or reused from the authors' own previous work); systematic reviews/meta-analyses *only* if they include primary data | Systematic reviews and meta-analyses without any primary data | Prevent duplication of previously synthesised results while retaining studies contributing original data. |
| 3. Microbial community focus | Analyses of bacterial gut microbiome communities containing ≥3 species across ≥2 genera | Studies focusing solely on non-bacterial communities (e.g., virome, mycobiome, protozoa, metazoan) or analysing <3 bacterial species | Ensure inclusion of studies assessing complex bacterial communities relevant to host–microbiota interactions. |
| 4. Anatomical focus | Analyses of the fore-gut or hind-gut microbiome of primates | Studies on non-gut microbiomes (e.g., skin, oral, genital) or on gut microbiomes of non-primate taxa | Focus on gut-associated bacterial communities of primates. |
| 5. Study setting and population | Primates living in the wild, zoos, or sanctuaries; mixed studies where only wild/zoo data were extracted | Studies conducted exclusively on primates housed in laboratories or research facilities | Maintain conservation relevance and exclude primarily biomedical research contexts. |
| 6. Language | Publications in English | Publications in languages other than English | Ensure clarity and consistency in data interpretation and extraction. |

(i) Dissertations and theses were excluded because their content is likely to overlap with articles subsequently published in peer-reviewed journals (Astiazarán-Azcárraga et al., 2024).

(ii) Systematic reviews and meta-analyses that did not include primary data were excluded to avoid duplication of previously synthesised results. However, such articles were included if they presented primary data (i.e., newly generated data or data reused from the author's previous work). In these cases, both the primary and secondary data were extracted for the host species included in the study.

(iii) Articles that did not analyse gut bacterial microbiome communities were excluded. The key criterion was a focus on bacteria; therefore, studies exclusively examining other microbial components of the gut microbiome, such as the virome (viruses), mycobiome (fungi), protozoa, or metazoans, were excluded. The second key criterion was a focus on microbial communities. For this review, a bacterial community was defined as comprising at least three species across a minimum of two genera; studies focusing on fewer were excluded. This threshold was applied to refine the review to studies explicitly investigating bacterial community structure and function; those that facilitate broader inferences to be made about the relationship between primate hosts and their gut microbiota.

(iv) Articles that did not analyse the foregut- or hindgut-microbiomes of primate, specifically, were excluded. Accordingly, studies focusing on gut microbiomes of non-primate host taxa were excluded, as were studies examining other body site microbiomes (e.g., skin or oral cavity microbiomes).

(v) Studies conducted exclusively on primates housed in laboratory or research facility settings were excluded to maintain a clear conservation focus. While some laboratory-based studies address relevant biological questions, their primary focus is typically biomedical and therefore falls outside the scope of this review. Studies on primates housed in zoos or sanctuaries were retained, as these often aimed to investigate gut bacterial microbiomes in relation to behaviour, diet, and/or conservation. In cases where samples were collected from primates in both wild or zoo settings and laboratory or research facilities, only data from wild or zoo-housed individuals were extracted.

(vi) Studies published in languages other than English were excluded to ensure accuracy and consistency in data extraction and interpretation.

Table SIII: The categories and respective data points extracted from each article.

| DATA EXTRACTED | DESCRIPTION | LABEL IN SUPPLEMENTARY FILE 1 |
|---|---|---|
| Publication ID | A unique numerical identifier (1–261) was assigned to each publication based on the chronological order of data extraction. | Publication_ID |
| Publication authors | Author names were extracted in the form of in-text citations (e.g., Bloggs et al.). When multiple articles shared the same first author name, a distinguishing lowercase letter was appended to each (e.g., Bloggs et al. (a), Bloggs et al. (b), irrespective of publication year, in chronological order of data extraction. The lowercase letter was added irrespective of publication year to aid data analysis. Consequently, these lowercase letters can be seen in the publication authors data point column in Supplementary File 1. | Authors |
| Year of publication | The year in which each article was published was recorded from the article metadata or header information. | Publication_year |
| Title of publication | The full title of each publication was recorded from the article metadata. | Publication_title |
| Journal title | The name of the journal in which each article was published was recorded. | Journal |
| Host common name | A common name was extracted for each host primate species studied. The common name of a single species often varied between articles and in its IUCN listing, therefore a single standardised name was selected and applied consistently across all records for that species. | Host_common_name |

| | | |
|---|---|---|
| Host species (IUCN-recognised) | Primate taxonomy has changed over time, so the binomial nomenclature reported in each article was cross-checked against the IUCN Red List. The IUCN-recognised binomial name was extracted in all cases to ensure consistency. | Host_species_IUCN |
| Host species (in article) | The binomial name used in each article was recorded to maintain transparency and allow readers to trace the original taxonomy in the supplementary material. | Host_species_publication |
| Host family (IUCN-recognised) | The taxonomic family of each host species, as recognised by the IUCN rather than the original article, was extracted in all cases to ensure consistency. | Host_family_IUCN |
| Host genus (IUCN-recognised) | The taxonomic genus of each host species, as recognised by the IUCN rather than the original article, was extracted in all cases to ensure consistency. | Host_genus_IUCN |
| Conservation status | The conservation status of each host species, as recognised by the IUCN rather than the original article, was extracted in all cases to ensure consistency and to reflect the species' current threat status. | Conservation_status_IUCN |
| Citation for conservation status | The citation listed on each host species' IUCN Red List page was recorded to allow readers to refer back to the original source material if needed. | Citation_for_conservation_status |
| Country of sample origin | The country of origin for each primate faecal sample was extracted based on the article's description. | Country_of_sample_origin |
| Region of sample origin | The region (continent) of origin for each primate faecal sample was extracted based on the article's description. | Region_of_sampling |
| Site name | The name of each site where samples were collected was | Wild_or_captive_site |

| | | |
|---|---|---|
| | extracted from the article's description. When multiple articles referred to the same site using different names or spelling variations, a single name was chosen and used consistently to standardise the data for analysis. | |
| Lifestyle type | The environmental setting of the sampled primates was extracted (e.g., wild, captive, unclear). | Lifestyle_type |
| Coordinates | If GPS coordinates were explicitly reported, these were used directly. When coordinates were not provided, we approximated them using the site name via Google Maps (2025). For studies that only specified the country of sample origin, we used the coordinates of the country's capital city to enable a general spatial overview and highlight potential geographic research gaps. All coordinates were converted to latitude and longitude using the WGS 84 datum. | Coordinates |
| Research scope | The study aim(s) of each article were extracted and grouped into one of 15 defined scopes (Supplementary File 3) to facilitate synthesis and enable more effective analysis of the dataset. | Scope |
| Sample type | The method used to collect sample material was extracted for each study (e.g., faecal samples, rectal samples). | Sample_type |
| Total samples analysed | The number of samples analysed was extracted from each article. When only the number of samples collected was reported, it was assumed that all collected samples were analysed. | Total_samples |
| Total individual hosts | The number of individual hosts from which samples were analysed was extracted. If only the number of hosts | Total_individuals |

| | | |
|---|---|---|
| | sampled was reported, it was assumed all sampled individuals were analysed. | |
| Preservation method | The method(s) used to preserve samples prior to freezing were extracted from each article. Where variations in terminology occurred for the same preservation method, terms were standardised. These methods were then grouped into eight categories to facilitate synthesis. | Preservation_meth_prior_freeze |
| Sequencing technology | The sequencing technology used in each article was extracted (e.g., Illumina, 454 (Roche), PacBio). | Sequencing_technology |
| Sequencing Generation | The generation of the sequencer(s) used in each article was extracted (e.g. First Generation, Second Generation). | Seq_gen |
| Next Generation Sequencing approach | The NGS approach used in each article was extracted (e.g., 16S rRNA amplicon sequencing, shotgun metagenomics sequencing). | NGS_approach |
| Amplicon marker gene | If 16S rRNA amplicon sequencing was used as the NGS approach, the targeted amplicon marker gene was extracted when reported. | Amplicon_marker |

Table SIV: Trait dataset used to assess influence on species study count.

| TRAIT | DESCRIPTION | SOURCE |
|---|---|---|
| Family | Species family according to IUCN. | IUCN, 2025a |
| Genus | Species genus according to IUCN. | IUCN, 2025a |
| Species | Species name according to IUCN. | IUCN, 2025a |
| Studied | Binominal: yes (1) or no (0) if species had been studied. | Supplementary File 1 |
| Study_count | Review species study count | Supplementary File 1 |

| | | |
|---|---|---|
| ResearchLegacy_genus | Total number of studies retrieved on Web of Science for each primate genera. | Web of Science, 2026 |
| BodyMass_kg | Mean body mass (kg), compiled from multiple published sources and made publicly available in primate traits dataset. | Galán-Acedo et al., 2019 |
| Locomotion | Locomotion (arboreal, terrestrial, both), compiled from multiple published sources and made publicly available in primate traits dataset. | Galán-Acedo et al., 2019 |
| DietAct | Diel activity (diurnal, nocturnal, cathemeral), compiled from multiple published sources and made publicly available in primate traits dataset. | Galán-Acedo et al., 2019 |
| TrophicGuild | Trophic guild (dietary type e.g. frugivore, folivore), compiled from multiple published sources and made publicly available in primate traits dataset. | Galán-Acedo et al., 2019 |
| Habitat_Forest | Binominal: yes (1) or no (0) if species has been recorded inhabiting this habitat type. | Galán-Acedo et al., 2019 |
| Habitat_Savana | Binominal: yes (1) or no (0) if species has been recorded inhabiting this habitat type. | Galán-Acedo et al., 2019 |
| Habitat_Shrubland | Binominal: yes (1) or no (0) if species has been recorded inhabiting this habitat type. | Galán-Acedo et al., 2019 |
| Habitat_Grassland | Binominal: yes (1) or no (0) if species has been recorded inhabiting this habitat type. | Galán-Acedo et al., 2019 |
| Habitat_Wetlands | Binominal: yes (1) or no (0) if species has been recorded inhabiting this habitat type. | Galán-Acedo et al., 2019 |
| Habitat_Rocky_areas | Binominal: yes (1) or no (0) if species has been recorded inhabiting this habitat type. | Galán-Acedo et al., 2019 |
| Habitat_Desert | Binominal: yes (1) or no (0) if species has been recorded inhabiting this habitat type. | Galán-Acedo et al., 2019 |

| | | |
|---|---|---|
| Habitat_Count | A count of the number of different habitat types that the species was recorded inhabiting, as recorded in a publicly available primate traits dataset. | Galán-Acedo et al., 2019 |
| HomeRange_ha | Species home range size (ha), compiled from multiple published sources and represented as a mean and made publicly available in primate traits dataset. | Galán-Acedo et al., 2019 |
| IUCN_status | The IUCN status of each species. | IUCN, 2025a |
| Pop_T | The IUCN population trend of each species, as recorded in a publicly available primate traits dataset. | Galán-Acedo et al., 2019 |
| Africa | Binominal: yes (1) or no (0) if species has been recorded inhabiting this region. | IUCN, 2025a |
| Asia | Binominal: yes (1) or no (0) if species has been recorded inhabiting this region. | IUCN, 2025a |
| North_America | Binominal: yes (1) or no (0) if species has been recorded inhabiting this region. | IUCN, 2025a |
| South_America | Binominal: yes (1) or no (0) if species has been recorded inhabiting this region. | IUCN, 2025a |
| Europe | Binominal: yes (1) or no (0) if species has been recorded inhabiting this region. | IUCN, 2025a |
| Oceania | Binominal: yes (1) or no (0) if species has been recorded inhabiting this region. | IUCN, 2025a |
| Continent_Count | A count of the number of regions that each species is distributed within. | IUCN, 2025a |

### Data analysis

*Primate taxonomic coverage*

Initially, data were manually curated in Excel. Taxonomic data were extracted for all primates studied and cross-checked against the IUCN Red List (IUCN, 2025a). Updated

IUCN binomial nomenclature was applied to ensure consistency with current species recognition. Across all articles, primate taxonomic information was successfully retrieved at the family level. Genus-level information could not be retrieved for two taxa, and species-level information could not be retrieved for 17 taxa (including the two taxa lacking genus-level identification). Taxa for which taxonomic identification could not be resolved were excluded from downstream analyses. Difficulties in taxonomic identification most commonly arose when primates were reported using non-specific labels (e.g. "*Cebus* sp." or *"Loris*"), with additional searches of supplementary materials failing to resolve these identifications.

Next, analyses were conducted in R (version 4.4.3; RStudio version 2026.04.0+526). Research effort was quantified separately at the species, genus, and family levels in R by counting each taxon once per publication. For this, the *Publication_ID*, *Host_species_IUCN*, *Host_genus_IUCN*, and the *Host_family_IUCN* data points were used (see Table SIII for descriptions). Hybrid taxa were retained if both parental species were reported but excluded if not. Summary statistics, including the mean and range of studies, were calculated across all taxa at the species, genus, and family levels. A stacked bar plot was created in R to illustrate the proportion of species studied within each primate family. The plot used data compiled from the IUCN Red List, including family names and total species counts, combined with our dataset recording the number of species studied per family.

A summary table was generated in R to report the number and proportion of species ever studied in the wild, captivity, or both contexts. Using the *Publication_ID*, *Host_species_IUCN*, and *Lifestyle_type* data points, each species was counted once per lifestyle type to calculate total counts and percentages of primate species studied in each setting.

Pie charts were created using Canva (https://www.canva.com/) to illustrate the taxonomic and conservation status coverage of primates studied in gut bacterial microbiome research. The proportions used in these pie charts were calculated using the *Publication_ID*, *Host_species_IUCN*, *Host_genus_IUCN*, *Host_family_IUCN*, and *Conservation_status_IUCN* data points in R.

Additionally, to support our discussion of the taxonomic coverage and geographic distribution, we compiled the current list of primate species, genera, and families from the IUCN Red List (IUCN, 2025a) and cross-checked it against our curated dataset. For each species, data were extracted of its region(s) of distribution, whether it is a multi-regional species, whether it is native or introduced to a region, and whether it has been sampled globally, within region, and within region in the wild (Supplementary File 2).

*Geographic distribution of sampling*

A global sampling heatmap was created in R (packages: *dplyr* (Wickham et al., 2023a), *tidyr* (Wickham et al., 2024a), *sf* (Pebesma, 2018; Pebesma & Bivand, 2023), *rnaturalearth* (Massicotte & South, 2023), *ggplot2* (Wickham, 2016), and *ggnewscale* (Campitelli, 2025)) to illustrate the global research distribution and effort. Countries were colour-graded by publication frequency, and field site points scaled by the number of studies and coloured by lifestyle type of the host inhabiting the site (wild, captive, unclear). This map was created using the *Publication_ID*, *Country_of_sample_origin*, *Lifestyle_type*, *Wild_or_captive_site*, and *Coordinates* data points. The number of distinct publications per country was summarised to assess research distribution globally, and coordinates were used to assess the number of studies per field site. GPS coordinates were used when they were explicitly reported in the article. If GPS data were not provided, the site name in combination with Google Maps (2025) were used to approximate coordinates. For studies that only specified the country of sample origin, and no other location information, we defaulted to the coordinates of the country's capital city to allow for a general spatial overview of sampling distribution and highlight potential geographic research gaps.

Bar plots were created in R (packages: *dplyr*, *ggplot2, scales* (Wickham et al., 2023b), *patchwork* (Pedersen, 2024), *RColorBrewer* (Neuwirth, 2022)) to illustrate the geographic patterns of research effort across regions based on the country and region of sample origins, using the *Publication_ID*, *Country_of_sample_origin*, and *Region_of_sampling* data points. A stacked bar plot (a) shows the proportional contribution of each country to the total number of studies within each region. For this, each study-country combination within a region was counted once, to ensure that multi-country studies did not inflate region study counts, and country counts that contributed less than 5% to a region's total were grouped as "Other" for clarity within the figure. A horizontal bar plot (b) shows the proportion of studies that included each region.

A global primate species richness map was created in R (packages: *terra* (Hijmans, 2025), *ggplot2*, *sf*, and *grDevices* (R Core Team, 2025)) using raster data downloaded from Biodiversitymapping.org (Jenkins et al., 2013; Pimm et al., 2014; Biodiversitymapping, n.d.). The raster was projected to WGS84 and converted to a dataframe for plotting. The map was visualised using a continuous colour gradient from blue (low richness) to red (high richness), overlaid on a world map. This map was created to visualise global primate species richness and to aid comparison with the global distribution of geographic research and sampling effort.

*Temporal trends in publication frequency, research scope, and methodologies employed*

Temporal trends in publication output were assessed using a stacked bar chart produced in R (packages: *dplyr*, *ggplot2*, *readr* (Wickham et al., 2024b)). Distinct publications were identified using *Publication_ID*, and annual publication counts were summarised by grouping publications by *Publication_year* and sequencing generation (*Seq_gen*). Counts were visualised as stacked bars to illustrate changes in research output and sequencing methodology over time.

Journal count and diversity were assessed using the *Publication_ID* and *Journal* data points, by counting the number of unique publication identifiers within each distinct journal. These data were also filtered to included only articles published between 2020 and 2025 to provide an insight into which journals are on the forefront of this research field.

Research scopes were derived from the study aims of each of the included studies and manually organised into 15 categories (the research scopes); studies with aims fitting multiple categories were assigned to each relevant scope (Supplementary File 3). The temporal distribution of research scopes was visualised using a bubble plot (R packages: *dplyr*, *ggplot2*) using the *Publication_ID*, *Publication_year*, and *Scope* data points to assess how research focus and effort has shifted over time. Distinct publication-scope combinations were identified and summarised to determine the count of studies per scope per year. Summary statistics were also calculated (range, mean, median, standard deviation, and interquartile range; including only years with at least one study published) to support the visual patterns (R package: *dplyr*).

Descriptive statistics were calculated to summarise methodological aspects of the studies included in the systematic review. The number of samples analysed, and the number of individual hosts sampled per study, were analysed using the *Total_samples* and *Total_individuals* data points to calculate the mean, median, range, and standard deviation of each data point. The same approach was conducted for Sample type (*Sample_type*), preservation methods (*Preservation_meth_prior_freeze*), sequencing technologies (*Sequencing_technology*), NGS approaches (*NGS_approach*), and amplicon markers (*Amplicon_marker*).

*GLMM and PGLMM*

To assess potential factors that predict species study count, we considered multiple factors, including species ecological traits, geographical distribution, and taxonomic information (Table SIV) (Supplementary File 5).

Two generalised linear mixed models (GLMMs) were created to account for missing information using the *glmmTMB* package (Brookes et al., 2017). GLMM1 included only categorical predictors (*Locomotion*, *DielAct*, *TrophicGuild*, *IUCN_status*, *Pop_T*) as fixed effects, with *Family* and *Genus* specified as random effects. All "NA" and "NI" values in categorical variables were recoded as "Unknown." GLMM2 extended this structure by incorporating both categorical predictors and scaled numerical variables (*Continent_Count*, *BodyMass_kg*, *Habitat_Count*, *ResearchLegacy_genus*) as fixed effects, while retaining *Family* and *Genus* as random effects. Before fitting GLMM2, any species with missing values in numerical predictors were excluded from the dataset (202 species).

To assess if primates have similar study effort after accounting for ecological and geographical traits, we used a phylogenetic GLMM (PGLMM) in the *phyr* package (Ives et al., 2025). Numerical predictors (*BodyMass_kg*, *HomeRange_ha*, *Habitat_Count*, *Continent_Count*) were used as fixed effects and *Species*, with covariance (phylogenetic correlation matrix), as a random effect.

### RESULTS AND DISCUSSION

#### Results and discussion: tables

Table SV: The number of publications per journal.

| JOURNAL | PUBLICATION COUNT |
| --- | --- |
| American Journal of Primatology | 31 |
| Scientific Reports | 20 |
| Frontiers in Microbiology | 17 |
| The ISME Journal | 10 |
| Microbial Ecology | 8 |
| Animal Microbiome | 7 |

| | |
|---|---|
| Animals | 7 |
| Journal of Medical Primatology | 6 |
| PLoS One | 6 |
| Ecology and Evolution | 5 |
| Global Ecology and Conservation | 5 |
| Microbiome | 5 |
| Molecular Ecology | 5 |
| Primates | 5 |
| npj Biofilms and Microbiomes | 5 |
| American Journal of Physical Anthropology | 4 |
| BMC Genomics | 4 |
| International Journal of Primatology | 4 |
| Biology | 3 |
| FEMS Microbiology Ecology | 3 |
| Frontiers in Ecology and Evolution | 3 |
| Microbiology Spectrum | 3 |
| Msystems | 3 |
| Nature Communications | 3 |
| iScience | 3 |
| Cell Reports | 2 |
| Frontiers in Veterinary Science | 2 |
| Integrative Zoology | 2 |
| Journal of Wildlife Diseases | 2 |
| Microbiology | 2 |
| PeerJ | 2 |
| Proceedings of the Royal Society B-Biological Sciences | 2 |
| Science | 2 |
| eLife | 2 |
| mBio | 2 |
| mSphere | 2 |
| Acta Biologica Colombiana | 1 |
| Animal Behaviour | 1 |
| Applied Microbiology | 1 |

| | |
|---|---|
| Applied and Environmental Microbiology | 1 |
| Archives of Microbiology | 1 |
| Arquivo Brasileiro De Medicina Veterinaria E Zootecnia | 1 |
| BMC Ecology and Evolution | 1 |
| BMC Microbiology | 1 |
| BioMed Research International | 1 |
| Biology Letters | 1 |
| Biotropica | 1 |
| Brazilian Journal of Microbiology | 1 |
| Cell Host & Microbe | 1 |
| Communications Biology | 1 |
| Computational and Structural Biotechnology Journal | 1 |
| Current Biology | 1 |
| Current Microbiology | 1 |
| Current Research in Microbial Sciences | 1 |
| Diversity | 1 |
| EcoHealth | 1 |
| Ecotoxicology and Environmental Safety | 1 |
| Emerging Infectious Diseases | 1 |
| Environmental DNA | 1 |
| Environmental Microbiology | 1 |
| Environmental Microbiology Reports | 1 |
| Evolutionary Applications | 1 |
| FEMS Microbiology Letters | 1 |
| Folia Primatologica | 1 |
| Frontiers in Cellular and Infection Microbiology | 1 |
| Frontiers in Endocrinology | 1 |
| Frontiers in Microbiomes | 1 |
| Genes | 1 |
| Gut Microbes | 1 |
| Gut Pathogens | 1 |
| ISME Communications | 1 |
| Infection, Genetics and Evolution | 1 |

| | |
|---|---|
| Innovation | 1 |
| Integrative and Comparative Biology | 1 |
| International Journal for Parasitology: Parasites and Wildlife | 1 |
| Journal of Animal Ecology | 1 |
| Journal of Applied Animal Research | 1 |
| Journal of Applied Microbiology | 1 |
| Journal of Microbiological Methods | 1 |
| Journal of Microbiology | 1 |
| Journal of Veterinary Science | 1 |
| Journal of Zoological and Botanical Gardens | 1 |
| Malaysian Applied Biology | 1 |
| Microbial Ecology in Health and Disease | 1 |
| MicrobiologyOpen | 1 |
| Microorganisms | 1 |
| Molecular Biology and Evolution | 1 |
| Molecular Ecology Resources | 1 |
| Nature Microbiology | 1 |
| Oecologia | 1 |
| PLoS Pathogens | 1 |
| Pathogens | 1 |
| Proceedings of the National Academy of Sciences | 1 |
| Royal Society Open Science | 1 |
| Science Advances | 1 |
| Science of the Total Environment | 1 |
| Veterinary Medicine and Science | 1 |
| Veterinary World | 1 |
| Zoo Biology | 1 |
| iMetaOmics | 1 |

Table SVI: The number of studies of which focused on each research scope.

| SCOPE | NUMBER OF STUDIES |
|---|---|
| Diet-related Impacts on Microbiota | 77 |
| Wild Environment and Habitat Influences on Microbiota | 71 |
| Microbiota Characterisation and Diversity | 56 |
| Other Host-specific Factors | 53 |

| | |
|---|---|
| Evolution of Host Species and their Microbiota | 44 |
| Captive Environment Influences on Microbiota | 41 |
| Anthropogenic Alterations and Impacts on Microbiota | 35 |
| Pathogen Prevalence | 31 |
| Social and Behavioural Impacts on Microbiota | 30 |
| Microbiota Functionality | 25 |
| Antibiotic-related Impacts on Microbiota | 20 |
| Host Health Implications | 14 |
| Temporal Variation in the Microbiota | 7 |
| Methodological Factors on Microbiota | 5 |
| Conservation Applicability | 2 |

Table SVII: Examples of how microbiome research on primates has conservation relevance.

| **MICROBIOME INSIGHT** | **STUDY SNAPSHOT** | **CONSERVATION RELEVANCE** | **SPECIES (NOT ALL LISTED)** | **REFERENCE** |
|---|---|---|---|---|
| Revealing interactions between gut pathogens and other microbial communities | In wild mouse lemurs, associations were made between bacterial microbiota and certain parasites. For example *Eimeria* spp. impacted microbiota diversity, whilst cestodes interacted with several bacterial orders. This study highlights some of the complex dynamics within gut microbiome communities. | Gut microbiome composition can be shaped by parasitic infections, with potential consequences for host health. These interactions may disrupt key microbial functions related to digestion and immunity. Recognising and monitoring such dynamics is important for understanding disease risk and guiding conservation efforts in primate populations, particularly in environments where parasitic burden is high. | Rufous mouse lemur (*Microcebus rufus*) 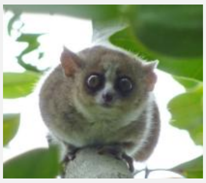 Microcebus_rufus_001.jpg: Photo: Alex Dunkel (Visionholder); Camera: Freddie Barberderivative work: WolfmanSF, CC BY-SA 3.0 <https://creativecommons.org/licenses/by-sa/3.0>, via Wikimedia Commons | Aivelo & Norberg, 2018 |
| Revealing effects of habitat degradation on gut microbiota | A study on Udzungwa red colobus monkeys found that individuals in | Primates have co-evolved with their gut microbiota to meet ecological | Udzungwa red colobus (*Piliocolobus gordonorum*) | Barelli et al., 2015 |

| | | | | |
|---|---|---|---|---|
| | disturbed forests had significantly lower gut microbiota diversity. Shifts in microbial composition likely reflected reduced dietary plant diversity, and functional analysis suggested a loss of microbes involved in detoxifying plant tannins. | challenges, including detoxifying plant compounds. Habitat degradation disrupts many environment–host–microbiome interactions, potentially leading to the loss of beneficial microbial functions. This study reveals how microbiome disruption reflects hidden host-health costs of ecosystem disturbance. | 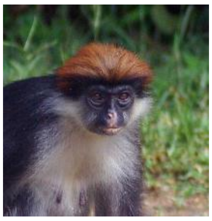 Stevage, CC BY-SA 3.0 <https://creativecommons.org/licenses/by-sa/3.0>, via Wikimedia Commons | |
| Identifying antibiotic resistance linked to human proximity | A study on wild and captive baboons found that those living closer to humans had altered microbiota composition and a broader range of antibiotic resistance genes. These findings suggest that contact with humans promotes alterations in the gut microbiome of non-human primates which lead to changes in the ecology of gut communities. | Antibiotic resistance is a global threat to wildlife, with the spread posing health risks for primates. This study shows how human encroachment and also captivity can expand the antibiotic “resistome" of primates, potentially compromising their ability to respond to infections. Managing the human to primate interface is critical for safeguarding primate health and limiting cross-species transmission of antibiotic resistant bacteria. | Kinda baboon (*Papio kindae*) 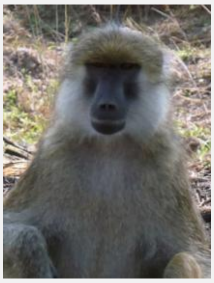 Kenneth Chiou, CC BY-SA 4.0 <https://creativecommons.org/licenses/by-sa/4.0>, via Wikimedia Commons | Tsukayama et al., 2018 |

| | | | | |
|---|---|---|---|---|
| Assessing the impact of human food consumption on gut microbiota | A study on wild Arunachal macaques found that their gut microbiota closely resembled that of humans, likely due to frequent consumption of human food waste near settlements. This raises concerns about reduced access to natural food resources, increased human–primate conflict, and potential exposure to human pathogens. | Human food waste provides easily accessible resources for omnivorous primates, highlighting their dietary plasticity. However, this reliance raises conservation concerns, including increased human–primate conflict, reduced access to natural food, and heightened risk of reverse zoonosis. Effective management should include limiting access to human waste, restoring natural foraging options, and educating the public on the mutual health risks of close interactions. | Arunachal macaque (*Macaca munzala*) 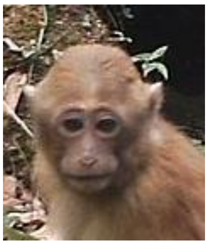 Nandini Velho, CC BY-SA 3.0 <https://creativecommons.org/licenses/by-sa/3.0>, via Wikimedia Commons | Ghosh et al., 2020 |
| Tracking seasonal shifts in the gut microbiome | A study on black howler monkeys found that seasonal dietary shifts were associated with shifts in gut microbiome composition and function. In periods of lower energy intake, microbiota enhanced energy production, buffering the host against nutritional stress. | Primate resilience and adaptability are challenged by many factors, especially increasing human impacts. Yet many primates face natural annual lean periods. This study highlights the microbiome's critical role in supporting primate host-health during these times, ultimately boosting | Yutacán black howler monkey (*Alouatta pigra*) 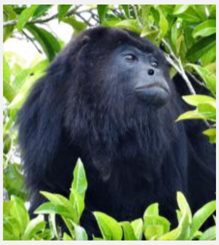 María Eugenia Mendiola González, CC BY-SA 4.0 <https://creativecommons.org/licenses/by-sa/4.0>, via Wikimedia Commons | Amato et al., 2015 |

| | | | | |
|---|---|---|---|---|
| | | survivability for these populations. | | |
| Assessing impacts of captivity on gut microbiota | A comparative study of five primate genera found that captivity altered the gut microbiome in all species, with the strongest effects in folivores. These included changes in diversity and reductions in fibre-degrading taxa. The authors recommended incorporating more natural browse into folivore diets. | Understanding how captivity alters the primate gut microbiome, especially in dietary specialists, can improve husbandry practices, reduce diet-related health issues, and support the success of long-term captive breeding programs. | Guereza (Colobus guereza) 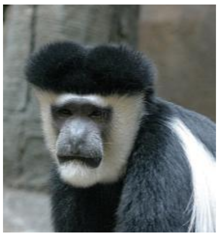 en:User:Cburnett, CC BY-SA 3.0 <http://creativecommons.org/licenses/by-sa/3.0/>, via Wikimedia Commons | Frankel et al., 2019 |
| Monitoring gut microbiome changes during reintroduction | A study on reintroduced woolly monkey's microbiota showed that diversity increased post-release, likely related to increased dietary diversity. Microbiome composition and functionality likewise increased post-release, suggesting a positive health impact of life in a more natural environmental setting. | Monitoring the gut microbiome during reintroduction provides valuable insight into how primates respond to release. Increased microbial diversity following reintroduction may reflect improved health from wild foraging. These findings can help guide pre-release management and support the long-term success of reintroduction programs. | Common woolly monkey (*Lagothrix lagothricha*) 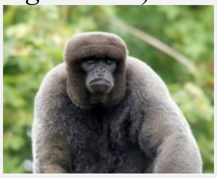 © Hans Hillewaert. <a title="© Hans Hillewaert" href="https://commons.wikimedia.org/wiki/File:Lagothrix_lagotricha_(male).jpg"></a> |

Table SVIII: The total number of studies that sampled species within each primate family.

| **HOST FAMILY** | **NUMBER OF STUDIES** |
|---|---|
| Cercopithecidae | 127 |
| Hominidae | 52 |
| Lemuridae | 42 |
| Atelidae | 31 |
| Indriidae | 30 |
| Hylobatidae | 21 |
| Callitrichidae | 18 |
| Cheirogaleidae | 15 |
| Cebidae | 11 |
| Lorisidae | 9 |
| Pitheciidae | 7 |
| Lepilemuridae | 5 |
| Daubentoniidae | 3 |
| Galagidae | 3 |
| Aotidae | 2 |

Table SIX: The total number of studies that sampled species within each primate genera.

| **HOST GENUS** | **NUMBER OF STUDIES** |
|---|---|
| *Macaca* | 43 |
| *Rhinopithecus* | 40 |
| *Gorilla* | 35 |
| *Pan* | 34 |
| *Alouatta* | 29 |
| *Propithecus* | 28 |
| *Lemur* | 26 |
| *Papio* | 25 |
| *Eulemur* | 24 |
| *Colobus* | 21 |
| *Cercopithecus* | 19 |
| *Trachypithecus* | 17 |
| *Ateles* | 14 |
| *Piliocolobus* | 13 |
| *Microcebus* | 12 |
| *Theropithecus* | 11 |
| *Chlorocebus* | 10 |
| *Pygathrix* | 10 |
| *Varecia* | 10 |
| *Hylobates* | 9 |
| *Nomascus* | 9 |

| | |
|---|---|
| *Callithrix* | 8 |
| *Cebus* | 8 |
| *Lagothrix* | 8 |
| *Leontopithecus* | 8 |
| *Pithecia* | 7 |
| *Symphalangus* | 7 |
| *Callimico* | 6 |
| *Cercocebus* | 6 |
| *Cheirogaleus* | 6 |
| *Hoolock* | 6 |
| *Indri* | 6 |
| *Nycticebus* | 6 |
| *Tamarinus* | 6 |
| *Lepilemur* | 5 |
| *Oedipomidas* | 5 |
| *Pongo* | 5 |
| *Semnopithecus* | 5 |
| *Nasalis* | 4 |
| *Sapajus* | 4 |
| *Avahi* | 3 |
| *Daubentonia* | 3 |
| *Erythrocebus* | 3 |
| *Hapalemur* | 3 |
| *Leontocebus* | 3 |
| *Xanthonycticebus* | 3 |
| *Aotus* | 2 |
| *Cebuella* | 2 |
| *Galago* | 2 |
| *Mandrillus* | 2 |
| *Mico* | 2 |
| *Otolemur* | 2 |
| *Plecturocebus* | 2 |
| *Prolemur* | 2 |
| *Saguinus* | 2 |
| *Saimiri* | 2 |
| *Brachyteles* | 1 |
| *Cacajao* | 1 |
| *Miopithecus* | 1 |
| *Mirza* | 1 |
| *Phaner* | 1 |
| *Presbytis* | 1 |
| *Procolobus* | 1 |

Table SX: The total number of studies that sampled each primate species.

| HOST SPECIES | NUMBER OF STUDIES |
|---|---|
| *Gorilla gorilla* | 33 |

| | |
|---|---|
| *Pan troglodytes* | 32 |
| *Lemur catta* | 26 |
| *Rhinopithecus roxellana* | 26 |
| *Macaca mulatta* | 21 |
| *Alouatta pigra* | 20 |
| *Eulemur rubriventer* | 17 |
| *Propithecus verreauxi* | 17 |
| *Rhinopithecus bieti* | 15 |
| *Colobus guereza* | 14 |
| *Papio hamadryas* | 13 |
| *Alouatta palliata* | 12 |
| *Trachypithecus francoisi* | 12 |
| *Cercopithecus ascanius* | 11 |
| *Papio anubis* | 11 |
| *Theropithecus gelada* | 11 |
| *Alouatta caraya* | 10 |
| *Alouatta seniculus* | 10 |
| *Papio cynocephalus* | 10 |
| *Propithecus coquereli* | 10 |
| *Ateles belzebuth* | 9 |
| *Pygathrix nemaeus* | 9 |
| *Rhinopithecus brelichi* | 9 |
| *Varecia variegata* | 9 |
| *Ateles hybridus* | 8 |
| *Lagothrix lagothricha* | 8 |
| *Macaca fuscata* | 8 |
| *Piliocolobus badius* | 8 |
| *Pan paniscus* | 7 |
| *Callimico goeldii* | 6 |
| *Cebus capucinus* | 6 |
| *Cercopithecus neglectus* | 6 |
| *Eulemur fulvus* | 6 |
| *Eulemur rufifrons* | 6 |
| *Indri indri* | 6 |
| *Pithecia pithecia* | 6 |
| *Propithecus diadema* | 6 |
| *Symphalangus syndactylus* | 6 |
| *Tamarinus imperator* | 6 |
| *Ateles geoffroyi* | 5 |
| *Callithrix geoffroyi* | 5 |
| *Callithrix jacchus* | 5 |
| *Chlorocebus aethiops* | 5 |
| *Chlorocebus sabaeus* | 5 |
| *Eulemur macaco* | 5 |
| *Gorilla beringei* | 5 |
| *Hylobates lar* | 5 |
| *Leontopithecus rosalia* | 5 |
| *Nomascus hainanus* | 5 |

| | |
|---|---|
| *Ateles fusciceps* | 4 |
| *Cercopithecus diana* | 4 |
| *Chlorocebus pygerythrus* | 4 |
| *Colobus angolensis* | 4 |
| *Colobus polykomos* | 4 |
| *Hoolock tianxing* | 4 |
| *Macaca fascicularis* | 4 |
| *Macaca thibetana* | 4 |
| *Microcebus rufus* | 4 |
| *Nasalis larvatus* | 4 |
| *Nycticebus bengalensis* | 4 |
| *Propithecus edwardsi* | 4 |
| *Trachypithecus cristatus* | 4 |
| *Cercocebus chrysogaster* | 3 |
| *Cercopithecus cephus* | 3 |
| *Cercopithecus wolfi* | 3 |
| *Cheirogaleus medius* | 3 |
| *Colobus vellerosus* | 3 |
| *Daubentonia madagascariensis* | 3 |
| *Erythrocebus patas* | 3 |
| *Eulemur coronatus* | 3 |
| *Eulemur flavifrons* | 3 |
| *Eulemur mongoz* | 3 |
| *Eulemur rufus* | 3 |
| *Microcebus griseorufus* | 3 |
| *Microcebus murinus* | 3 |
| *Nomascus annamensis* | 3 |
| *Nomascus leucogenys* | 3 |
| *Oedipomidas geoffroyi* | 3 |
| *Oedipomidas oedipus* | 3 |
| *Piliocolobus gordonorum* | 3 |
| *Pongo abelii* | 3 |
| *Pongo pygmaeus* | 3 |
| *Sapajus apella* | 3 |
| *Semnopithecus vetulus* | 3 |
| *Trachypithecus leucocephalus* | 3 |
| *Varecia rubra* | 3 |
| *Xanthonycticebus pygmaeus* | 3 |
| *Avahi laniger* | 2 |
| *Callithrix penicillata* | 2 |
| *Cebuella pygmaea* | 2 |
| *Cercocebus agilis* | 2 |
| *Cercopithecus campbelli* | 2 |
| *Cercopithecus nictitans* | 2 |
| *Cercopithecus petaurista* | 2 |
| *Chlorocebus djamdjamensis* | 2 |
| *Eulemur albifrons* | 2 |
| *Hylobates pileatus* | 2 |

| | |
|---|---|
| *Leontocebus weddelli* | 2 |
| *Leontopithecus chrysomelas* | 2 |
| *Leontopithecus chrysopygus* | 2 |
| *Lepilemur mustelinus* | 2 |
| *Macaca arctoides* | 2 |
| *Macaca assamensis* | 2 |
| *Macaca nemestrina* | 2 |
| *Macaca silenus* | 2 |
| *Macaca sylvanus* | 2 |
| *Mico argentatus* | 2 |
| *Microcebus danfossi* | 2 |
| *Nomascus concolor* | 2 |
| *Otolemur crassicaudatus* | 2 |
| *Papio ursinus* | 2 |
| *Prolemur simus* | 2 |
| *Propithecus candidus* | 2 |
| *Propithecus tattersalli* | 2 |
| *Saguinus mystax* | 2 |
| *Saimiri boliviensis* | 2 |
| *Semnopithecus priam* | 2 |
| *Trachypithecus mauritius* | 2 |
| *Trachypithecus phayrei* | 2 |
| *Alouatta guariba* | 1 |
| *Aotus azarae* | 1 |
| *Aotus nigriceps* | 1 |
| *Ateles chamek* | 1 |
| *Avahi peyrierasi* | 1 |
| *Brachyteles hypoxanthus* | 1 |
| *Cacajao calvus* | 1 |
| *Callithrix aurita* | 1 |
| *Callithrix jacchus x Callithrix penicillata* | 1 |
| *Callithrix penicillata x Callithrix geoffroyi* | 1 |
| *Cebus imitator* | 1 |
| *Cercocebus atys* | 1 |
| *Cercopithecus hamlyni* | 1 |
| *Cercopithecus mitis* | 1 |
| *Cercopithecus mona* | 1 |
| *Cercopithecus roloway* | 1 |
| *Cheirogaleus crossleyi* | 1 |
| *Eulemur collaris* | 1 |
| *Galago moholi* | 1 |
| *Galago senegalensis* | 1 |
| *Hapalemur aureus* | 1 |
| *Hapalemur griseus* | 1 |
| *Hoolock leuconedys* | 1 |
| *Hylobates abbotti* | 1 |
| *Hylobates agilis* | 1 |
| *Lagothrix flavicauda* | 1 |

| | |
|---|---|
| *Leontocebus fuscicollis* | 1 |
| *Lepilemur grewcockorum* | 1 |
| *Lepilemur ruficaudatus* | 1 |
| *Lepilemur sahamalaza* | 1 |
| *Macaca mulatta x Macaca fascicularis* | 1 |
| *Macaca munzala* | 1 |
| *Macaca sinica* | 1 |
| *Mandrillus leucophaeus* | 1 |
| *Mandrillus sphinx* | 1 |
| *Microcebus lehilahytsara* | 1 |
| *Miopithecus ogouensis* | 1 |
| *Mirza coquereli* | 1 |
| *Nomascus gabriellae* | 1 |
| *Nycticebus coucang* | 1 |
| *Nycticebus javanicus* | 1 |
| *Nycticebus menagensis* | 1 |
| *Otolemur garnettii* | 1 |
| *Papio kindae* | 1 |
| *Papio kindae x Papio ursinus* | 1 |
| *Papio papio* | 1 |
| *Phaner pallescens* | 1 |
| *Piliocolobus rufomitratus* | 1 |
| *Piliocolobus tephrosceles* | 1 |
| *Plecturocebus cupreus* | 1 |
| *Plecturocebus moloch* | 1 |
| *Plecturocebus oenanthe* | 1 |
| *Pongo pygmaeus x Pongo abelii* | 1 |
| *Presbytis femoralis* | 1 |
| *Presbytis robinsoni* | 1 |
| *Presbytis siamensis* | 1 |
| *Procolobus verus* | 1 |
| *Pygathrix nigripes* | 1 |
| *Rhinopithecus avunculus* | 1 |
| *Saguinus midas* | 1 |
| *Saimiri cassiquiarensis* | 1 |
| *Sapajus nigritus* | 1 |
| *Semnopithecus entellus* | 1 |
| *Semnopithecus schistaceus* | 1 |
| *Tamarinus labiatus* | 1 |
| *Trachypithecus crepusculus* | 1 |
| *Trachypithecus obscurus* | 1 |

Table SXI: The number of studies that analysed samples collected from each country.

| COUNTRY | NUMBER OF STUDIES |
|---|---|
| China | 77 |
| Madagascar | 47 |

| | |
|---|---|
| United States of America | 34 |
| Mexico | 21 |
| Central African Rep. | 18 |
| Uganda | 18 |
| Ethiopia | 14 |
| Tanzania | 14 |
| Malaysia | 11 |
| Colombia | 10 |
| Costa Rica | 10 |
| Argentina | 9 |
| Brazil | 9 |
| Cameroon | 9 |
| Ecuador | 8 |
| Japan | 8 |
| Kenya | 8 |
| Nicaragua | 8 |
| Congo | 7 |
| Dem. Rep. Congo | 7 |
| Singapore | 6 |
| Vietnam | 6 |
| South Africa | 5 |
| Thailand | 5 |
| France | 4 |
| Ghana | 4 |
| United Kingdom | 4 |
| Czechia | 3 |
| Côte d'Ivoire | 3 |
| Germany | 3 |
| Namibia | 3 |
| St. Kitts and Nevis | 3 |
| Switzerland | 3 |
| Australia | 2 |
| Gabon | 2 |
| India | 2 |
| Peru | 2 |
| Sri Lanka | 2 |
| Algeria | 1 |
| Austria | 1 |
| Belgium | 1 |
| Bolivia | 1 |
| Denmark | 1 |
| Indonesia | 1 |
| Ireland | 1 |
| Netherlands | 1 |
| Senegal | 1 |
| Sierra Leone | 1 |
| Slovakia | 1 |
| Spain | 1 |

| | |
|---|---|
| Taiwan | 1 |
| Zambia | 1 |

Table SXII: The number, and proportion, of studies that used each type of preservation method.

| PRESERVATION METHOD | STUDY COUNT | PERCENTAGE |
|---|---|---|
| Cold storage methods | 93 | 35.63 |
| Stabilising buffers | 82 | 31.42 |
| Ethanol | 57 | 21.84 |
| Transport media | 13 | 4.98 |
| Lysis buffers | 7 | 2.68 |
| FTA cards | 5 | 1.92 |
| Drying methods | 4 | 1.53 |
| Fixative buffers | 2 | 0.77 |

Table SXIII: The number, and proportion, of studies that used each sequencing technology.

| SEQUENCING TECHNOLOGY | STUDY COUNT | PERCENTAGE |
|---|---|---|
| Illumina | 204 | 78.16 |
| 454 (Roche) | 17 | 6.51 |
| Not used | 14 | 5.36 |
| PacBio | 6 | 2.30 |
| Sanger | 5 | 1.92 |
| Ion Torrent (PGM) | 2 | 0.77 |
| MGI Tech | 2 | 0.77 |
| Oxford Nanopore Technologies | 2 | 0.77 |

Table SXIV: GLMM1 coefficients.

| VARIABLE | ESTIMATE | STD. ERROR | Z VALUE | PR(>\|Z\|) |
|---|---|---|---|---|
| (Intercept) | 0.175772 | 0.836711 | 0.210074 | 0.83361 |
| LocomotionBOTH | 0.691225 | 0.356648 | 1.938117 | 0.052609 |
| LocomotionT | 0.767465 | 0.439093 | 1.74784 | 0.080492 |
| LocomotionUnknown | 0.071448 | 1.989494 | 0.035913 | 0.971352 |
| DielActD | 0.40738 | 0.782812 | 0.520406 | 0.60278 |
| DielActN | -1.1224 | 0.82952 | -1.35307 | 0.176033 |
| DielActUnknown | 2.600528 | 2.198092 | 1.183084 | 0.236776 |
| TrophicGuildFolivore_frugivore | 0.51714 | 0.34497 | 1.499086 | 0.133851 |
| TrophicGuildFrugivore | -0.57469 | 0.375329 | -1.53117 | 0.125727 |
| TrophicGuildGummivore | 0.179009 | 0.731999 | 0.244548 | 0.806806 |
| TrophicGuildInsectivore | -1.06894 | 1.014357 | -1.05381 | 0.291972 |
| TrophicGuildOmnivore | -0.46072 | 0.407434 | -1.13078 | 0.258149 |
| TrophicGuildUnknown | -2.19676 | 0.636333 | -3.45222 | 0.000556 |
| IUCN_statusDD | -18.322 | 3966.691 | -0.00462 | 0.996315 |
| IUCN_statusEN | 0.17774 | 0.296829 | 0.598796 | 0.549309 |
| IUCN_statusLC | -0.02528 | 0.401946 | -0.0629 | 0.94985 |
| IUCN_statusNE | -20.7262 | 5352.498 | -0.00387 | 0.99691 |

| | | | | |
|---|---|---|---|---|
| IUCN_statusNT | 0.052287 | 0.46429 | 0.112616 | 0.910335 |
| IUCN_statusUnknown | -5.35309 | 1.244552 | -4.30122 | 1.70E-05 |
| IUCN_statusVU | 0.005555 | 0.339009 | 0.016386 | 0.986927 |
| Pop_TI | 1.7685 | 0.931133 | 1.8993 | 0.057525 |
| Pop_TS | 0.784509 | 0.415241 | 1.889289 | 0.058853 |
| Pop_TUnknown | 0.690798 | 0.320055 | 2.158372 | 0.030899 |

Table SXV: GLMM2 coefficients.

| VARIABLE | ESTIMATE | STD. ERROR | Z VALUE | PR(>\|Z\|) |
|---|---|---|---|---|
| (Intercept) | 0.522665 | 0.800767 | 0.652705 | 0.513946 |
| LocomotionBOTH | 0.490195 | 0.387535 | 1.264905 | 0.205905 |
| LocomotionT | 0.394134 | 0.476726 | 0.826752 | 0.408377 |
| LocomotionUnknown | -21.442 | 27511.74 | -0.00078 | 0.999378 |
| DielActD | 0.178272 | 0.763751 | 0.233417 | 0.815438 |
| DielActN | -0.91262 | 0.773574 | -1.17975 | 0.2381 |
| DielActUnknown | -17.1662 | 5038.519 | -0.00341 | 0.997282 |
| TrophicGuildFolivore_frugivore | 0.303849 | 0.359041 | 0.84628 | 0.397396 |
| TrophicGuildFrugivore | -0.78297 | 0.392151 | -1.9966 | 0.045869 |
| TrophicGuildGummivore | -0.1085 | 0.734735 | -0.14767 | 0.882606 |
| TrophicGuildInsectivore | -1.32723 | 0.960114 | -1.38237 | 0.166858 |
| TrophicGuildOmnivore | -0.66562 | 0.434548 | -1.53175 | 0.125585 |
| TrophicGuildUnknown | -25.4932 | 142531.7 | -0.00018 | 0.999857 |
| IUCN_statusDD | -22.5193 | 40755.88 | -0.00055 | 0.999559 |
| IUCN_statusEN | 0.266345 | 0.304352 | 0.875123 | 0.381507 |
| IUCN_statusLC | 0.152642 | 0.420116 | 0.363332 | 0.716357 |
| IUCN_statusNE | -18.1597 | 7603.479 | -0.00239 | 0.998094 |
| IUCN_statusNT | 0.169042 | 0.499052 | 0.338725 | 0.734817 |
| IUCN_statusVU | 0.046292 | 0.345214 | 0.134096 | 0.893326 |
| Pop_TI | 1.146548 | 0.898164 | 1.276546 | 0.201762 |
| Pop_TS | 0.730308 | 0.410207 | 1.780342 | 0.07502 |
| Pop_TUnknown | 0.289032 | 0.353442 | 0.817762 | 0.413493 |
| Continent_Count_z | 0.208539 | 0.0878 | 2.375159 | 0.017541 |
| BodyMass_kg_z | 0.266875 | 0.131606 | 2.027837 | 0.042577 |
| Habitat_Count_z | 0.068881 | 0.115096 | 0.598466 | 0.549529 |
| HomeRange_ha_z | -0.07278 | 0.128615 | -0.56588 | 0.571477 |
| ResearchLegacy_genus_z | 0.064336 | 0.192399 | 0.334386 | 0.738088 |

Table SXVI: PGLMM coefficients.

| VARIABLE | ESTIMATE | STD_ERROR | Z_VALUE | P_VALUE |
|---|---|---|---|---|
| (Intercept) | -1.24717 | 0.60229 | -2.0707 | 0.038385 |
| BodyMass_kg | 0.02525 | 0.008296 | 3.0436 | 0.002338 |
| HomeRange_ha | -6.56E-05 | 0.000104 | -0.6328 | 0.526852 |
| Habitat_Count | 0.213203 | 0.146097 | 1.4593 | 0.144477 |
| Continent_Count | 1.096419 | 0.408439 | 2.6844 | 0.007266 |

### Results and discussion: figures

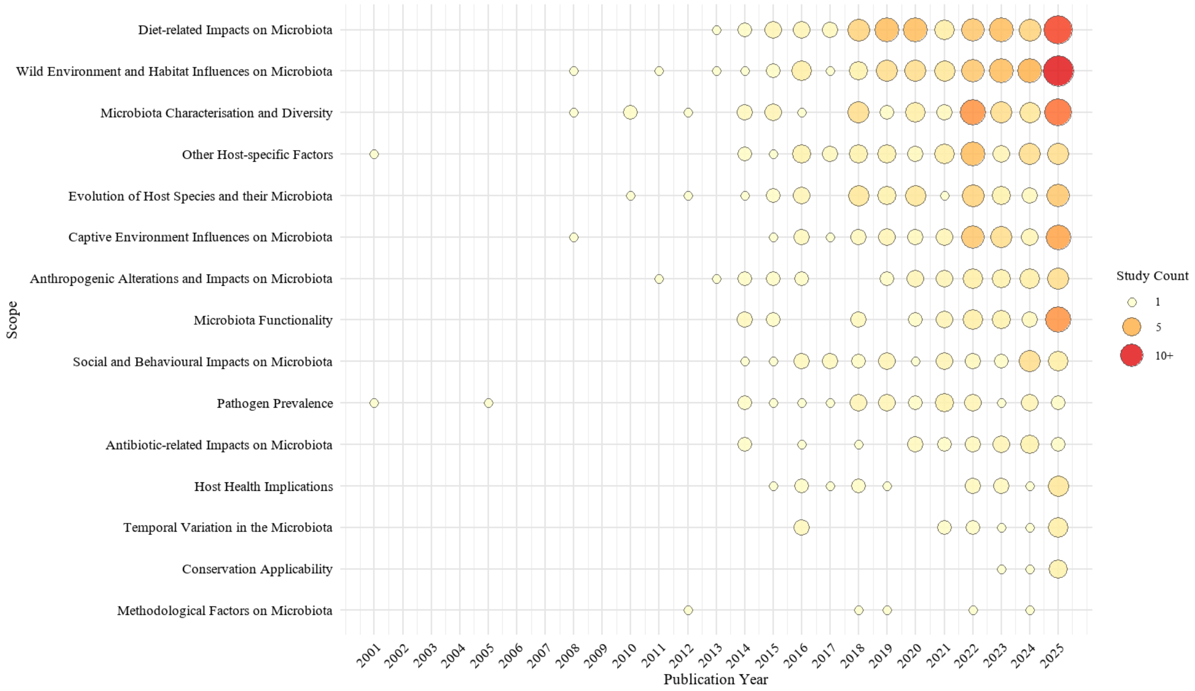


Figure S1: Bubble plot illustrates the frequency of different research scopes in primate microbiome studies from 2001-2025. The size of each bubble represents the number of studies focusing on a specific scope each year, with larger bubbles indicating more frequent studies. The colour gradient, from yellow to red, reflects the increasing frequency of studies, with yellow representing fewer studies and red indicating higher study counts.

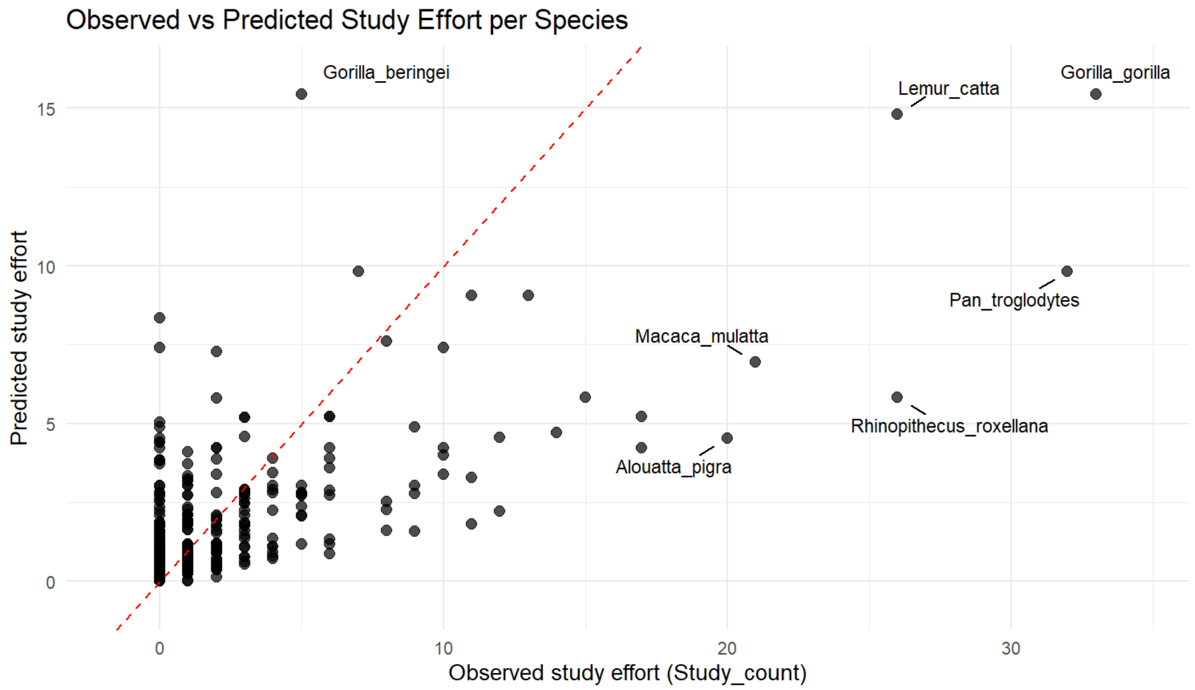


Figure S2: Observed versus predicted study count per primate species from GLMM1, which included categorical traits with missing values coded as “Unknown”. The dashed red line indicates perfect agreement between observed and predicted values. Deviations from this line highlight species that are over- or under-represented relative to model expectations.

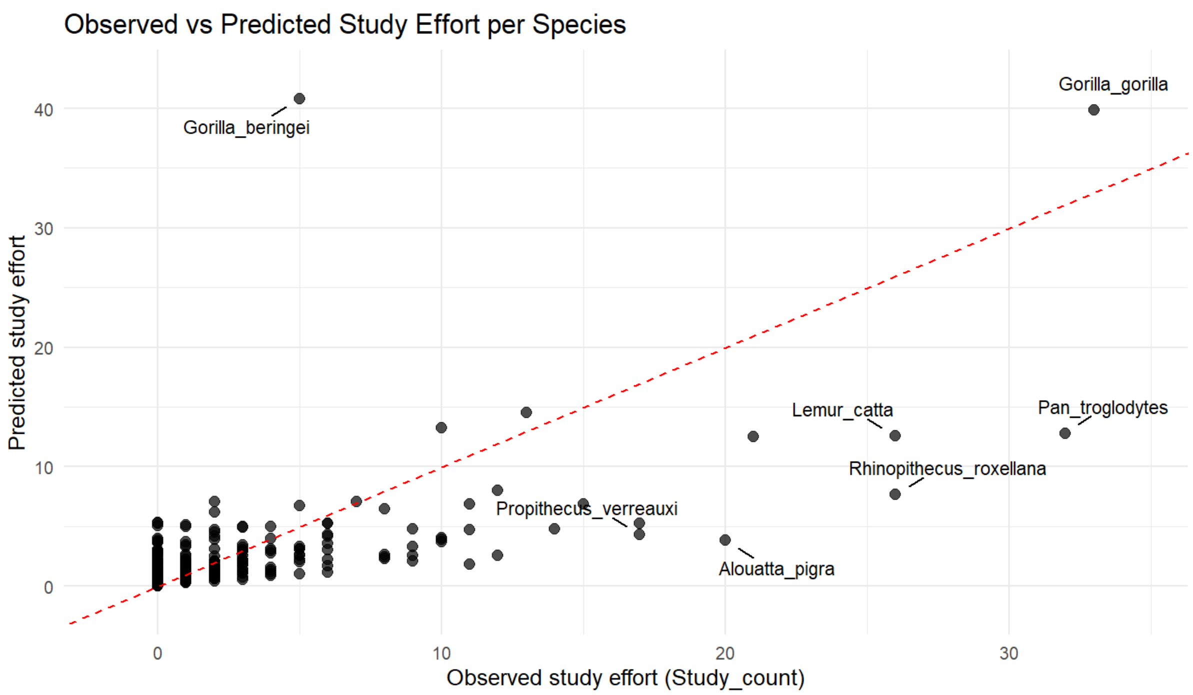


Figure S3: Observed versus predicted study effort per primate species from GLMM2, which only included species with complete numerical and categorical data. The dashed red line indicates perfect agreement between observed and predicted values. Points above the line indicate overprediction, while points below indicate underprediction. Deviations from this line highlight species that are over- or under-represented relative to model expectations.

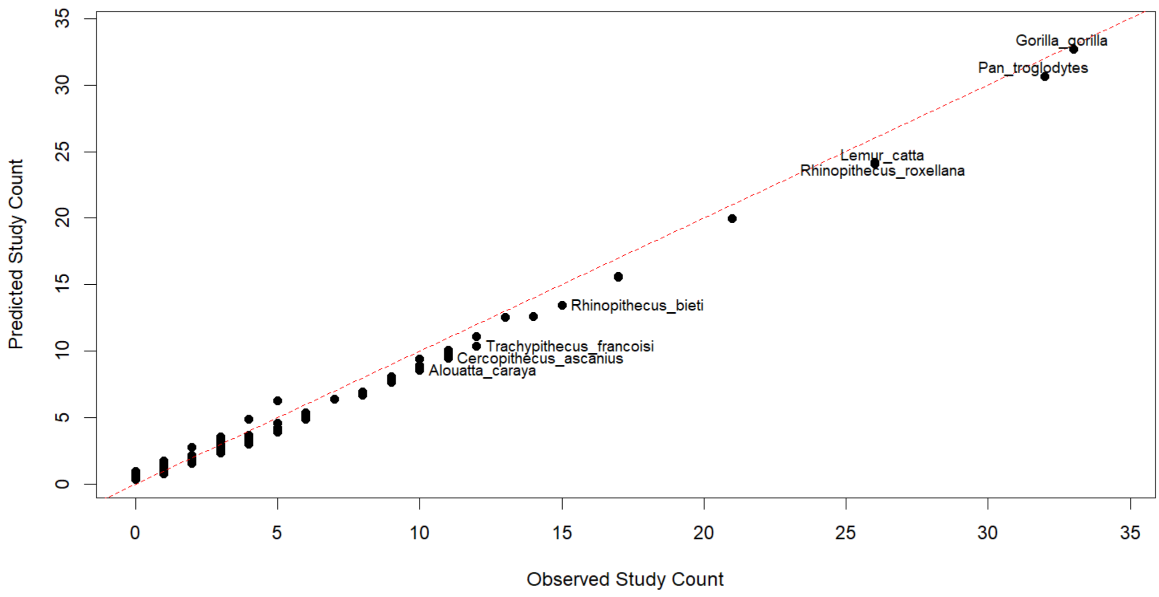


Figure S4: Observed versus predicted study effort per primate species from the PGLMM. The model incorporated phylogenetic relationships among species and only included species with complete numerical data. The dashed red line indicates perfect agreement between observed and predicted values. Points above the line indicate overprediction, while points below indicate underprediction. Deviations from this line highlight species that are over- or under-represented relative to model expectations.

**Detailed analysis by continent and respective countries on the taxonomic coverage and geographic distribution of primate bacterial microbiome studies**

*Asia: geographic distribution of sampling*

In Asia, primates are distributed across three regions: East Asia, South and Southeast Asia, and West and Central Asia (IUCN, 2025b). Species richness in Asia is highest in South and Southeast Asia (Figure 6). Our results show that Asia was the region with the highest proportion of studies (Figure 5a), spanning 10 countries (Table SXI). Within these, primates were most frequently sampled in China (77 studies), Malaysia (11 studies), and Japan (8 studies) (Figure 5b). China is an important country for primate conservation, as it hosts the highest primate species diversity in the northern hemisphere, many of these species are critically endangered, and many populations represent the most northern distribution of a species, genus, or family (Fan & Ma, 2018; Li et al., 2024). The most sampled site within Asia was the Beijing Zoo in China (13 studies), while the most sampled wild site was also in China; Shennongjia National Park (8 studies). While China currently receives most of the research effort in Asia, other countries such as Malaysia and Indonesia remain relatively undersampled (4.21%, n = 11; 0.38%, n = 1) despite substantial primate species richness of their own, with 26 and 65 extant species, respectively.

In other Asian countries, Indonesia is perhaps the greatest research void, as another high-priority area for primate conservation (Estrada et al., 2017). Our results show that only one study collected samples from primates in Indonesia (Cabana et al., 2019). This study analysed samples collected from wild Javan slow lorises (*Nycticebus javanicus*), captive greater slow lorises (*Nycticebus coucang*), and captive Philippine slow lorises (*Nycticebus menagensis*). Overall, only ~1.5% of Indonesia's wild primate species have been sampled. This is also somewhat surprising considering that Indonesia was the second most common location for primate conservation-related publications between 1994 and 2019 (de Figueiredo Machado et al., 2023). Furthermore, a review that looked at studies focusing on zoonotic pathogens (protozoa, GI parasites, viruses, bacteria, fungi, and blood-borne parasites) conducted between 1965 and 2023 on primates in Asia found that Indonesia was the joint top with Thailand (25 studies each) (Patouillat et al., 2024). Considering that methodologies to investigate zoonotic pathogens are somewhat similar, it suggests that barriers for research are dissimilar to those faced in the DRC, and that opportunities are there to expand this field of research towards Indonesian primates.

Another of the most-well studied countries by study count was Singapore (although there was a huge decrease in count following on from China). Since all the studies in Singapore were conducted on captive primates at the Singapore Zoo, the studies do not offer conservation value specifically to the extant species of Singapore; none of the country's species have been studied thus far. Several Asian countries with extant primate species have yet to be sampled for gut bacterial microbiome studies. These include Myanmar, Lao People's Democratic Republic (Lao) (both 20 sp.), Cambodia (14 sp.), Brunei Darussalam (11 sp.), Bangladesh (10 sp.), Bhutan (8 sp.), Nepal (4 sp.), Philippines, Pakistan (both 3 sp.), Hong Kong, Afghanistan (both 2 sp.), Timor-Leste, Saudi Arabia, and Yemen (all 1 sp.). Primates of Myanmar, Laos, and Cambodia are perhaps the greatest research voids other than Indonesia in Asia due to their high species richness.

*Asia: taxonomic coverage*

Concerning the taxonomic coverage, of the five primate families native to Asia, four have been represented in global bacterial microbiome research (80%). The only Asian family not represented was the Tarsiidae (tarsiers); which are the only family neither represented in this region nor studied in any context across the dataset. Species from the other four of these families (Cercopithecidae, Hylobatidae, Lorisidae, Hominidae) have been studied within Asia (either in the wild or captivity). Except for Hominidae (in Asia, the orangutans), members of the other three families have been sampled in the wild within Asia. Of the 19 genera currently recognised as native to Asia, 15 have been studied in bacterial microbiome research globally (78.95%). None of the following genera have been studied in the wild or captivity: *Loris* (Lorisidae); *Carlito*, *Cephalopachus*, *Tarsius* (Tarsiidae). All 15 studied genera include species that have been sampled in Asia (wild or captive); though only 12 have been sampled in the wild. The genera that have yet to be sampled in the wild are *Hylobates*, *Papio*, and *Pongo*. One genus that comprises multiple species (*Xanthonycticebus*) have only been represented by a single species, leaving the intra-genus diversity relatively unexplored.

At the species level, 51 of the currently recognised 130 Asian primate species have been included in global gut bacterial microbiome studies (39.23%). 46 of these species have been sampled within Asia (in either wild or captive settings) and five species have only been sampled outside of Asian countries (*Hoolock leuconedys, Macaca Silenus, Nomascus gabriellae, Pongo abelii, Semnopithecus entellus*). Of the 46 species sampled in Asia, 33 have been sampled in the wild. In Asia, two species are listed as Data Deficient on the IUCN Red List: the Bicoloured banded langur (*Presbytis bicolor*) found in Sumatra, Indonesia, alongside the Lariang tarsier (*Tarsius lariang*) from Sulawesi, Indonesia (IUCN, 2025a), investigating their gut bacterial microbiome could provide important data for these data devoid species which could potentially be used to support future IUCN Red List assessments.

*Africa: geographic distribution of sampling*

In Africa, primates are distributed across three regions: North Africa, sub-Saharan Africa, and Madagascar (IUCN, 2025b). Species richness is highest in Sub-Saharan Africa and Madagascar (Figure 6). Our results show that samples were collected from primates within Africa second-most frequently (Figure 5a), with 18 countries represented in our dataset (Table SXI). Among these, primates were most frequently sampled in Madagascar (47 studies), Uganda and the Central African Republic (CAR) (both 18) (Figure 5b). Across all regions, primates in Madagascar were the second most frequently sampled, only behind those in China. This aligns with a review by de Figueiredo Machado et al. (2023), which reported that Madagascar was the most common location for primate conservation-related publications between 1994-2019. Madagascar has been identified by Estrada et al. (2017) as one of four high-priority areas for primate conservation. Madagascar's primates span five families and are all endemic to the island (Ganzhorn et al., 1999); unfortunately, they are highly threatened by extinction (Michielsen et al., 2023). Our results show that, of all wild sites globally, three of the top six were in Madagascar: Ranomafana National Park (18 studies), Kirindy Forest, and Beza Mahafaly Special Reserve (both 11). Notably, Ranomafana National Park and Kirindy Forest were also two of the top four sites for data collected and published on primate field studies between 2011 to 2015 (Bezanson & McNamara, 2019).

As mentioned, primates of Uganda and the CAR have also been frequently sampled. In the CAR, the Dzanga-Sangha Protected Areas (DSPA) is a key field site, with 14 studies conducted there. The DSPA is home to two great ape species, the chimpanzee (*Pan troglodytes*) and the western lowland gorilla (*Gorilla gorilla gorilla*), as well as nine monkey species (Blom et al., 2005). The DRC is another high priority for primate conservation by Estrada et al. (2017). Unfortunately, only seven studies collected samples from primates in the DRC, all focusing on bonobos (*Pan paniscus*) (Moeller et al., 2016; Gaulke et al., 2018; Hickmott et al., 2022; Sanders et al., 2023; Rühlemann et al., 2024; Gao et al., 2025; Torfs et al., 2025). While studying the microbiome of this endangered species is clearly important, it represents just one of the country's fifty primate species, meaning that just 2% of the DRC's primates have been sampled in the wild. Countries in the eastern Congo Basin, such as the DRC, face logistical issues, armed conflict, and political instability, which hinder ecological research efforts (van der Hoek et al., 2021). Greater accessibility (roads, cities, urban areas) may also play a role in the choice of field site (Ellison et al., 2021). Several African countries with extant primate species have yet to be sampled for gut bacterial microbiome studies. These include Nigeria (27 species), Equatorial Guinea (24 species), Angola (23 species), Rwanda (18 sp.), Guinea (16 sp.), Burundi (15 sp.), Benin, Guinea-Bissau, Liberia, South Sudan, Togo (all 13 sp.), Burkina Faso, Mozambique, Somalia (all 9 sp.), Gambia, Malawi, Mali, Sudan (all 7 sp.), Zimbabwe (6 sp.), Chad (5 sp.), Eritrea, Eswatini, Mauritania, Niger (all 4 sp.), Botswana (3 sp.), Djibouti, Lesotho (both 2 sp.), Cabo Verde, Comoros, Egypt, Mauritius, Mayotte, Sao Tome and Principe, Morocco, Tunisia (all 1 sp.). Countries such as Nigeria, Equatorial Guinea, Angola, and Rwanda have very high primate species richness; yet little, if nothing, is currently known about their bacterial gut microbiota, highlighting large gaps in African primate bacterial microbiome research.

*Africa: taxonomic coverage*

Across Africa, 8 of 9 native primate families (88.9%) appear in microbiome studies; Lorisidae is the only African family absent, and Daubentoniidae has been studied only in captivity outside Africa, whereas the other seven families include wild-sampled species within Africa. Of the 42 genera currently recognised as native to Africa, 30 have been studied in bacterial microbiome research globally (71.43% ); however, four genera (*Daubentonia*, *Hapalemur*, *Mandrillus*, *Miopithecus*) are represented only by captive samples outside Africa, and several multi-species genera (e.g., *Erythrocebus*, *Miopithecus*, *Mirza*, *Phaner*) are each represented by just a single species. At the species level, 40.93% (88/215) of African species were studied, with 21 species only sampled outside Africa. The coverage is high at the family level, but decreases at the species level, highlighting notable gaps in understudied species, with limited intra-genus representation.

*South America: geographic distribution of sampling*

South America remains undersampled when compared to Asia and Africa (i.e., ~7.5% vs ~36% and ~34% of studies, Figure 5a), although it harbours very high primate species richness (Figure 6). The distribution of studies across South American countries was relatively even compared to African and Asian distributions (Figure 5b), with six countries sampled, most notably in Colombia (10 studies), Brazil and Argentina (9 studies each), and Ecuador (8 studies). One of these countries, Brazil, is another of the four high-priority areas for primate conservation listed by Estrada et al. (2017). Primate species richness is high in

Brazil, especially in the Amazon Basin (Figure 6). The total number of microbiome studies that collected samples in Brazil was low overall (3.45% of all publications) but moderate in terms of the region's total studies (39.13%). Like Indonesia, Brazil also ranked high in the review by de Figueiredo Machado et al. (2023) on primate conservation-related publications conducted between 1994 and 2019. Figueiredo Machado et al. reported that primates that exhibit both arboreal and terrestrial locomotion are more frequently studied than those that exhibit just one type or the other, and Patouillat et al. (2024) reported that semi-terrestrial species were more studied than strictly arboreal species. Most Brazilian primates are arboreal; only the capuchins (*Sapajus* spp. and *Cebus* spp.) are semi-terrestrial. It can be difficult to collect samples from arboreal primates due to difficulties locating them in the canopy, especially in dense forests. Even if defecation is observed, the sample can be easily lost by the time it reaches the forest floor. Nonetheless, compared to countries in other regions, the research effort is low. Increasing the overall research effort in Brazil and other South American countries is key for increasing our understanding of global primate gut bacterial microbiome biodiversity. Several South American countries with extant primate species have yet to be sampled for gut bacterial microbiome studies. These include Venezuela (22 sp.), French Guiana, Guyana (both 10 sp.), Suriname (9 sp.), Paraguay (5 sp.), and Uruguay (1 sp.).

*South America: taxonomic coverage*

Of the five primate families native to South America, all five have been represented in global bacterial microbiome research. Moreover, species from all five families have been studied within South America (either in the wild or captivity). Apart from the Pitheciidae, species from the remaining four families have been sampled in the wild within South America. Of the 24 genera currently recognised as native to South America, 20 have been studied in gut bacterial microbiome research globally (83.33%). None of the following genera have been studied in the wild or captivity: *Callicebus*, *Cheracebus, Chiropotes* (Pitheciidae); *Callibella* (Callitrichidae). Among the 20 genera studied, 15 include species that have been sampled in South America (wild or captive); 10 of which have been sampled in the wild. The genera that have yet to be sampled in the wild in South America are *Cacajao, Callibella, Callicebus, Callimico, Cebuella, Cebus, Cheracebus, Chiropotes, Oediopomidas, Mico, Plecturocebus, Pithecia, Saguinus,* and *Saimiri*. Several genera that comprise multiple species (*Brachyteles, Cacajao, Cebuella, Cebus, Mico, Pithecia*) have only been represented by a single species, which, as described for the coverage within Africa and Asia, leaves the intra-genus diversity relatively unexplored. At the species level, 41 of the currently recognised 177 South American primate species have been included in global bacterial microbiome studies (23.16%). 27 of these species have been sampled within South America (in either wild or captive settings) and 14 species have only been sampled outside of South America. Of the 27 species sampled in South America, 17 have been sampled in the wild.

In South America, there are numerous New World monkeys, particularly from Brazil, that are Data Deficient on the IUCN Red List. These include five saki monkeys of the genus Pithecia: Gray's bald-faced saki (*Pithecia irrorata*) found in Brazil, Bolivia, and Peru; Cazuza's saki (*Pithecia cazuzai*), Pissinatti's saki (*Pithecia pissinatti*), and Vanzolini's saki (*Pithecia vanzolinii*), all from Brazil; and Isabel's saki (*Pithecia isabela*) from Peru. Also in Brazil are two titi monkeys, Milton's titi monkey (*Plecturocebus miltoni*) and Stephen Nash's titi monkey (*Plecturocebus stephennashi*). Colombia is home to Hernández-Camacho's night monkey (*Aotus jorgehernandezi*) (IUCN, 2025a). Like the Data Deficient primates of Asia, investigating the gut bacterial microbiomes of these species could provide valuable information to support future IUCN Red List assessments.

*North America: geographic distribution of sampling*

In this review, we grouped North America with Mesoamerica (Central America) and the Caribbean Islands and labelled this North America. In this region, primates are naturally distributed across Mesoamerica but are non-native to North America and the Caribbean. However, introduced populations are now found in the USA and some Caribbean islands. Our results showed that primates were sampled in North American countries in ~18.5% of studies (Figure 5a), and that five countries were sampled in (Table SXI). Of these, primates were most frequently sampled in the USA (34), Mexico (21), and then Costa Rica (10) (Figure 5b). Across the entire dataset, primates situated in the USA were the third most studied (number of studies) and were included in ~61% of the region's total studies (Figure 5b). As mentioned, primates are not native to the USA, though there are established populations of introduced squirrel monkeys (*Saimiri* sp.), green monkeys (*Chlorocebus sabaeus*), and rhesus macaques (*Macaca mulatta*) in Florida (Anderson et al., 2017, Williams et al., 2021). Across all North American sampling sites, the Duke Lemur Center, situated in the USA, was the site most studied (16 studies). Established in 1966, this site is valuable for strepsirrhine primate conservation, having recorded data on over 4,200 individuals across 40 different taxa (lemurs, lorises, and galagos) (Zehr et al., 2014). Some of the valuable research at this site includes comparing the gut bacterial microbiome of captive strepsirrhines to their wild counterparts in Madagascar (e.g., Greene et al., 2019; Bornbusch et al., 2022).

Of the countries that have native populations of wild primates, Mexico has received relatively high attention (fourth in publication frequency) and was included in 37.5% of the region's total (Figure 5b). Moreover, all of Mexico's primate species (3) have been studied at least once. One of the field sites in Mexico, Palenque National Park, was a common sampling site (12 studies), meaning that it was sampled in almost 57% of all studies that sampled in Mexico. This bias is not too surprising considering that Palenque National Park was the third most common site overall for published primate studies between 2011 and 2015 (Bezanson & McNamara, 2019). This site is also where the Yutacán black howler monkey (*Alouatta pigra*) has been extensively studied, primarily by Amato et al. (e.g., 2013; 2014; 2015; 2016; 2017a; 2017b). Although research voids need to be addressed, site and species biases are also highly valuable. Repeat study of a species' gut microbiome at a single site can provide a deeper understanding, beyond baseline characterisations of the gut bacterial microbiome of the host, which can broaden our understanding of the importance of the microbiome for the host. Similar to the coverage of Mexican primates, taxonomic coverage across North American countries is also relatively high. The introduced but wild living green monkeys of St. Kitts and Nevis Island (Saiyed et al., 2025) have been studied (Asangba et al., 2019; Gomez et al., 2019; Asabanga et al., 2022). Several North American countries with extant primate species have yet to be sampled for gut bacterial microbiome studies. These include Panama (8 sp.), Guatemala, Honduras (both 3 sp.), Belize, Trinidad and Tobago (both 2 sp.), El Salvador (1 sp.), Barbados, Grenada, and Puerto Rico (all 1 sp. (introduced)).

*North America: taxonomic coverage*

North America has four primate families, which are native (Callitrichidae, Cebidae, Aotidae, Atelidae) and one which has been introduced (Cercopithecidae). Of these five families, four have been represented in global bacterial microbiome research (80%). The only South American family not represented was the Aotidae (night monkeys), although South American aotids have been studied elsewhere. Species from the other four families have all been studied within North America (either in the wild or captivity). Species from Cercopithecidae, Cebidae, and Atelidae have been sampled in the wild in this region, whereas species from Callitrichidae and Aotidae have not. Of the 10 genera currently recognised as native to North America (six native and four introduced), eight have been studied in bacterial microbiome research globally (80%). Among those studied, five include species that have been sampled in North America (wild or captive) (*Alouatta, Ateles, Cebus, Chlorocebus* (introduced), and *Oedipomidas*. Apart from *Oedipomidas*, species from the other four genera have been sampled from the wild in this region. At the species level, 12 of the currently recognised 16 North American primate species (12 native and four introduced) have been included in global gut bacterial microbiome studies (75%). Seven of these species have been sampled within North America (in either wild or captive settings) and five species have only been sampled outside of North America. Of the seven species sampled in North America, six have been sampled in the wild. Only one of the four species that have been introduced to North America, *C. sabaeus*, has been studied in the region. This species has only been studied in captivity within its native region of Africa. The other three species are of African (*Cercopithecus mona*, *Erythrocebus patas*) and Asian (*M. mulatta*) descent. These three species have all been studied in the wild and in captivity in their native regions.

*Europe and Oceania: geographic distribution of sampling*

Europe and Oceania are also two regions without native extant primate species; however, there are some introduced populations. Our results showed that primates were sampled in Europe and Oceania in ~4% and ~0.5% of studies, respectively (Figure 5a). In Europe, primates were most frequently sampled within France and the UK (both 4), Czechia, Germany, and Switzerland (all 3), and then Austria, Belgium, Denmark, Ireland, the Netherlands, Slovakia, and Spain (all 1) (Table SXI). In Europe, due to the lack of wild primates, sampling has only occurred at captive sites. European zoos play a role in global wildlife conservation efforts, like those in the USA and elsewhere, they can be valuable to primate conservation efforts in their own right. Perhaps the most interesting research void in Europe is that of the introduced and wild living Barbary macaques (*Macaca sylvanus*) of Gibraltar. These primates live side-by-side with humans, making them an interesting subject of study to investigate how human activities shape primate gut microbiome or vice versa. These macaques could also be studied in comparison to the populations that are living in their native range of the Atlas Mountains of Morocco, Tunisia, and Algeria, to investigate the impact of historical wildlife relocations and subsequent native gut microbiota shifts.

The two studies that were conducted within Oceania sampled primates in Australia (Figure 5b). These studies studied the gut microbiomes of chimpanzees at Taronga Zoo (Willenborg et al., 2022) and golden lion tamarins (*Leontopithecus rosalia*) at Adelaide Zoo (Lawless et al., 2025). Across Oceania introduced populations of long-tailed macaques (*Macaca fascicularis*) can be found on the island of Ngeaur in the Republic of Palau (Wheatley, 2011) and in Papua New Guinea. Like the macaques in Gibraltar, an opportunity for study does exist for these introduced primates.

*Europe and Oceania: taxonomic coverage*

As described, Europe and Oceania have one species each that has been introduced to their regions. Europe has the Barbary macaque (*M. sylvanus*) (native to Africa), and Oceania has the long-tailed macaque (*M. fascicularis*) (native to Asia). Both species have been represented in global bacterial microbiome research. In Europe, *M. sylvanus* has been sampled from within captivity but not in the wild. In Oceania, *M. fascicularis* has not been sampled in any context.
